# Enhanced Viral Detection in Grapevine via Exome Depletion and Next-Generation Sequencing (NGS)

**DOI:** 10.64898/2026.04.16.718969

**Authors:** Raúl Andrés Cuello, Diego Zavallo, Pablo Vera, Agustín Sattler, Andrea Puebla, Humberto Debat, Sebastián Gómez Talquenca, Sebastián Asurmendi

## Abstract

Grapevine (*Vitis vinifera L.*) is highly prone to viral infections that pose a significant threat to global viticulture sustainability. Traditional detection methods, such as PCR and ELISA, are limited to well-known pathogens, highlighting the need for more comprehensive and unbiased approaches. Here, we present the development of a cost-effective viral enrichment system adapted to next-generation sequencing (NGS) for the detection and characterization of grapevine viruses. Our strategy leverages hybridization-based capture using biotin-labeled cDNA probes hereafter named “Chloro-Zero”) designed to selectively deplete highly abundant host transcripts particularly plastid and ribosomal RNAs while preserving viral RNA. Probe design was informed by transcriptomic analysis of *V. vinifera*. We evaluated different subtractor-to-target RNA ratios, observing a consistent reduction of host RNA and a moderate enrichment of viral sequences. NGS analysis revealed improved recovery of low-abundance viral transcripts, with coverage levels comparable, to a certain extent, to those obtained using previously available commercial kits, but at a significantly lower cost. Although variability in depletion efficiency was observed, the results demonstrate the potential of this scalable and locally adaptable protocol for virome profiling in grapevines. By addressing key limitations of current depletion methods, our approach facilitates the detection of emerging viral threats and supports the development of more effective certification programs and sustainable management practices. Ongoing improvements in probe design and bioinformatic workflows are expected to enhance performance, providing a robust platform for broader applications in plant virology.

## 1. INTRODUCTION

Grapevine (*Vitis vinifera* L.) is one of the most economically important fruit crops worldwide, significantly contributing to the agricultural sectors of numerous countries through the production of wine, table grapes, raisins, and related products (Estêvão et al., 2024). In Argentina, the viticulture industry plays a critical role, with more than 220,000 hectares cultivated, primarily in the Andean provinces of Mendoza and San Juan (Alonso et al., 2024). Given the perennial nature of grapevines and their vegetative propagation, these plants are particularly prone to systemic infections caused by viruses and viroids, which have a profound impact on vine health and productivity (Debat et al., 2019a; Fuchs, 2025). The dissemination of viral pathogens poses a significant threat to viticulture sustainability, as viruses reduce grape yield and quality, leading to substantial economic losses (Fuchs, 2025).

More than 100 viruses and several viroids have been identified in grapevine globally, including the Grapevine leafroll-associated virus (GLRaV), Grapevine fanleaf virus (GFLV), Grapevine virus A (GVA), and Grapevine Pinot gris virus (GPGV) (Fuchs, 2025). While traditional methods for virus detection, such as PCR and ELISA, have been used extensively, they are limited to the detection of known pathogens (Javaran et al., 2021). High-throughput sequencing (HTS) technologies have emerged as a robust alternative, allowing for the comprehensive profiling of the “virome”, the entire set of viral agents present in grapevine tissues (Hadidi et al., 2016; Adams et al., 2009). This approach is essential for the identification of emerging viral pathogens and the characterization of mixed infections, which are common in grapevines (Coetzee et al., 2010; Debat et al., 2019b).

Recent advances in next-generation sequencing (NGS) and third-generation sequencing (TGS) technologies, such as Oxford Nanopore sequencing, have further revolutionized the field of grapevine virology (Phannareth et al., 2021). These methods enable the simultaneous detection of viruses and viroids without prior knowledge of their genome sequences, thereby providing a comprehensive and unbiased analysis of the grapevine virome (Miljanić et al., 2022; Fall et al., 2020). The ability to directly sequence RNA molecules and detect viral RNA modifications offers new opportunities to study the interactions between grapevine hosts and their associated pathogens (Badial et al., 2018). Several studies have successfully demonstrated the power of these sequencing techniques in elucidating the complex interactions between viral agents in grapevine tissues (Guček et al., 2017; Al Rwahnih et al., 2009).

In addition to virus detection, advances in bioinformatics tools for the *de novo* assembly of viral genomes have significantly improved our ability to reconstruct complete viral genomes from HTS data (Soltani et al., 2021). Various assembly approaches have been employed to optimize the recovery of viral sequences from metagenomic datasets (Thapa et al., 2024). These tools are essential for identifying unknown viruses and characterizing viral variants, which can have implications for virus evolution and epidemiology (Hily et al., 2020). Studies of grapevine tissues from different geographical regions have revealed the presence of unique viral species and variants that were previously undetectable using classical approaches (Debat et al., 2019b; Villamor et al., 2019).

To address the persistent challenges in grapevine virus detection and respond to the needs of the wine industry, we developed a comprehensive, cost-effective, and efficient method for detecting grapevine viruses using NGS technologies (Di Gaspero et al., 2022). The proposed methodological framework uses hybridization-capture techniques to enrich viral RNA before sequencing (Maina et al., 2024). This enriches viral RNA, enhancing sequencing coverage and sensitivity (Miozzi et al., 2013).

The potential impact of this work extends beyond virus detection. By characterizing the grapevine virome, it becomes possible to track the evolution of viral pathogens, identify potential quarantine risks, and support certification programs for virus-free propagation material (Maliogka et al., 2018; Abrahamian et al., 2025). This is particularly relevant given the significant costs and regulatory hurdles associated with the international movement of grapevine plant material (Campos et al., 2021). Furthermore, the ability to identify viruses in asymptomatic plants enables more effective disease management, which is critical for maintaining sustainable viticulture practices (Fuchs et al., 2025).

In summary, we developed a comprehensive NGS-based method for the detection and characterization of grapevine viruses, using advanced molecular and bioinformatics tools. This work addresses key challenges in virus diagnostics and contributes to the broader understanding of grapevine virus epidemiology and control (Jeger, 2020). The resulting platform offers versatile applications, from strengthening certification systems to supporting healthier and more competitive grapevine propagation at both national and international levels.

## 2. MATERIALS AND METHODS

### 2.1. Sample Collection and Preparation

*Vitis vinifera plant tissues used in this study were obtained from the EEA INTA Mendoza collection. Four accessions, 171, CH40, MB21, and P1103 were selected as putative virus-free sources of RNA for probe template production. In addition, a fifth accession, carrying multiple viral infections and referred to as Torrontés, was included as a test sample.* Samples were obtained by tissue disruption and immediately were frozen in liquid nitrogen and stored at -80°C until processing. Frozen tissue was ground into a fine powder using a pre-chilled mortar and pestle.

### 2.2. RNA Extraction and probe synthesis

The RNA extraction from tissue samples was carried out using the RNAprep Pure Plant Plus Kit (Tiangen Biotech, Beijing, China). Once the RNA was obtained, the samples were quantified and used to synthesize biotin-labeled cDNA probes. For this purpose, Biotin-16-dUTP (Merck) was employed, replacing dTTP in DNA labeling reactions at a ratio of 35% Biotin-16-dUTP and 65% dTTP. The synthesis was performed following the protocol provided by manufacturer (Thermo Fisher, Carlsbad, CA) to synthesize the first strand of cDNA using M-MLV RT starting from 1 µg of total RNA and each corresponding oligonucleotide. The biotinylated RNA-cDNA hybrid was cleaned with RNase H using the protocol suggested by the manufacturer (NEB, Ipswich, MA).

### 2.3. Enrichment of RNA of interest through hybridization and streptavidin pull-down

Hybridization-based capture of plant RNA to enrich viral RNA (Figure 1) was performed using two components: the biotin-labeled probe (hereafter referred to as the subtractor) and the plant RNA targeted for enrichment (hereafter referred to as the subtracted). Biotinylated cDNA probes (calculated in approximately average length ∼300 nt) were hybridized to Vitis vinifera total RNA in 30 µl reaction volumes, using RNA:cDNA at different molar ratios (1:1 and 1:2 molar ratios corresponding to 2.27 pmol and 4.55 pmol of biotinylated cDNA, respectively). Hybridization was carried out in a thermal cycler with the following program: 95 °C for 1 min, 85 °C for 30 s, 25 °C for 1 min, followed by a hold at 4 °C.

**Figure 1.**
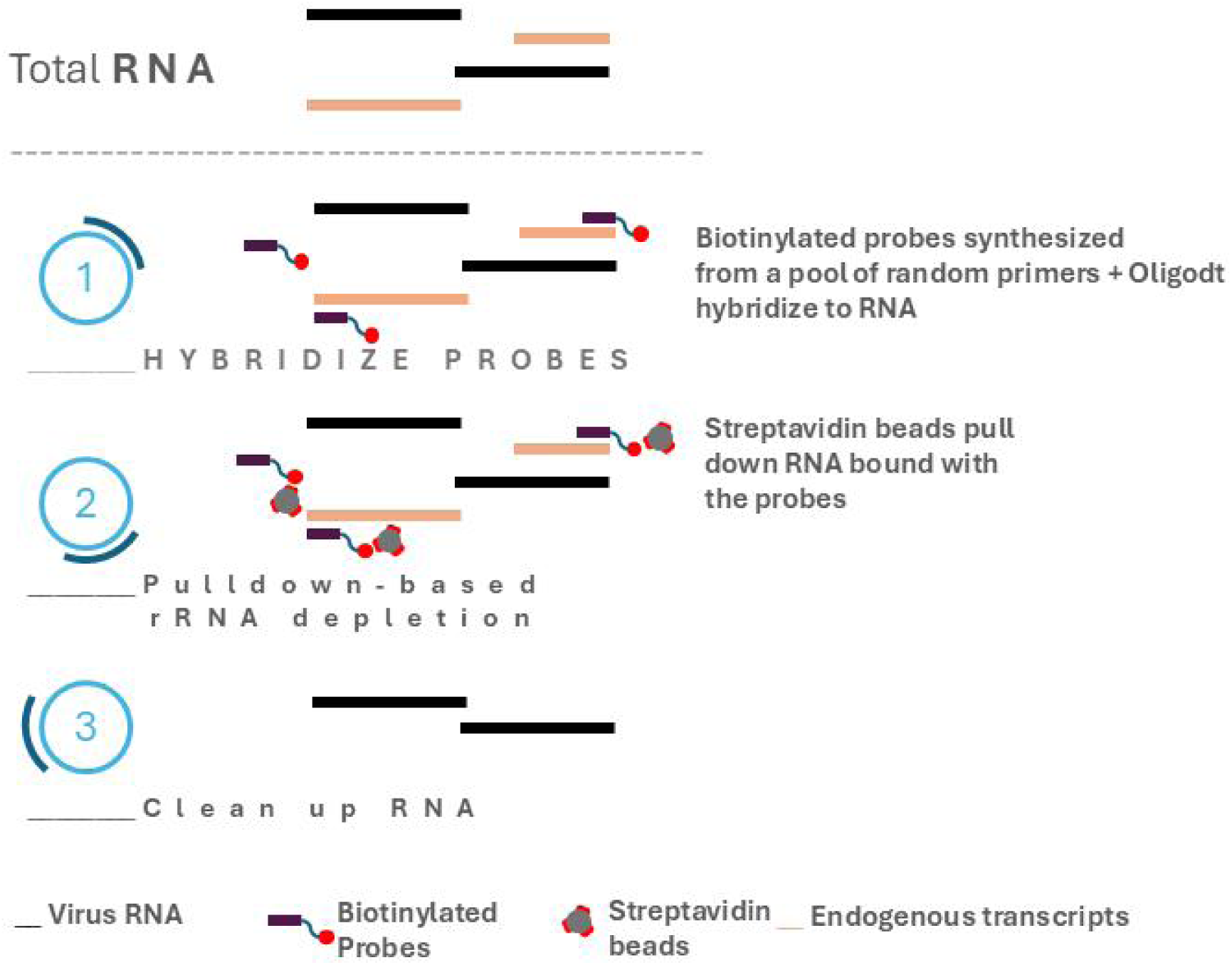
Conceptual diagram of expressed exome rejection. **Section 1**. Starting from our RNA extraction, biotin-labeled cDNA probes are made with a pool of random primers + oligo(dT). **Section 2**. Pull down of the hybrids formed between the labeled cDNA and the plant RNA using streptavidin beads. This captured exome is discarded, leaving the RNA that remains in suspension. **Section 3**. Once ready, the RNA is cleaned, to be used to evaluate the depth of viral enrichment through qPCR.

To capture RNA–cDNA hybrids, 60 µl of streptavidin-coated magnetic beads (10 mg/ml; Roche Diagnostics GmbH, Mannheim Germany) was added, providing approximately 1080 pmol of biotin-binding capacity per reaction (manufacturer specification: ≥1800 pmol biotin/mg × 0.60 mg beads). This represents a large molar excess of available binding sites relative to the amount of biotinylated probe, ensuring efficient hybrid recovery.

Before use, streptavidin-coated beads were pre-washed with TEN100 buffer (10 mM Tris-HCl pH 7.5, 1 mM EDTA, 1 M NaCl). After hybrid binding, bead–RNA complexes were placed on a magnetic stand to separate the beads, and washed with at least twice the original reaction volume of TEN100 buffer for 10 min at +15 to +25 °C. The complexes were again immobilized on a magnetic stand, and the resulting supernatant—containing the RNA depleted of complementary plant sequences (i.e., enriched viral RNA)—was collected and used for sequencing.

### 2.4. Improvement of probe synthesis (Chloro-Zero probes)

Considering the approach described by Forsythe et al. (2022), and to improve the exome-reduction system (Figure 2), specific primers (Supplementary Table 1A) were designed to target approximately 40% of the most highly expressed grapevine genes, based on our transcriptome dataset (Table 1 and Supplementary Table 2). This strategy aimed to objectively enhance the generation of biotin-labeled cDNA, in contrast to using random primers or oligo(dT). cDNA probe synthesis was performed using this primer set following the protocol described in Section 2.2.

**Figure 2.**
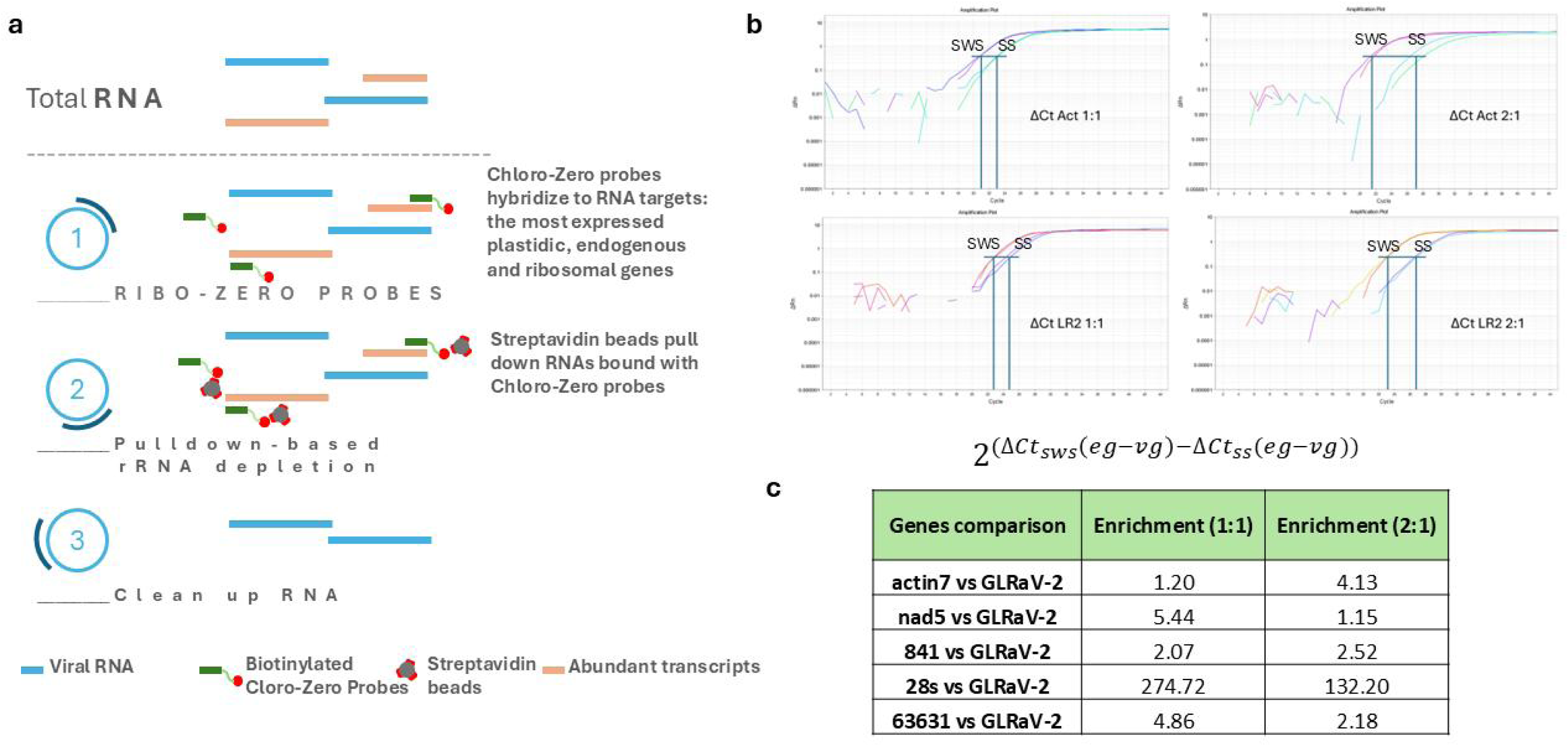
Conceptual diagram of accurate exome rejection with Chloro-Zero. 2a. Same methodology described in Figure 1 but with the replacement of the cDNA probes made from the primers batches (Chloro-Zero) listed in Table 1b. **2b.** Examples of real-time qPCR analyses of subtractions at different subtractor to target ratios, both 1:1 and 2:1, with the formula for calculating the associated ratios between subtracted RNA and non-subtracted RNA. ‘eg’ corresponds to the endogenous gene, ‘vg’ to the viral gene, ‘sws’ to the sample without subtraction, and ‘ss’ to the subtracted sample. **2c.** The table shows the differences corresponding to each enrichment rate, comparing some endogenous and ribosomal genes against a viral gene of GLRaV-2.

**Table 1.** Characterization of transcripts associated with plastids and ribosomes. 1a: Abbreviated list of the 200 most expressed genes in grapevine, representing a considerable percentage of reads (≈ 50% of the total reads) with their respective biological functions. **1b:** Chlro-Zero Batches of genes that target ribosomal, plastid, most expressed endogenous genes and some particular reference genes such as Actin and NAD. Primers used to target these genes are to improve the system for constructing labeled probes, replacing random primers and oligo(dT).

| Gene name | Counts | Description |
| --- | --- | --- |
| ViviCp001 | 23273776 | psbA photosystem II protein D1 |
| ViviM_p049 | 5864041 | psbA PSII 32 kDa protein |
| ViviCp029 | 3718155 | rbcL RuBisCO large subunit |
| ViviCp017 | 1760935 | psbC PSII 43 kDa protein |
| ViviM_p010 | 1692599 | rbcL ribulose-1,5-bisphosphate carboxylase/oxygenase large subunit |
| ViviCp016 | 815859 | psbD photosystem II protein D2 |
| ViviCp048 | 733230 | psbB photosystem II P680 chlorophyll A apoprotein |
| ViviCp052 | 418293 | petB cytochrome b6 |

b
| Batch 1 | Batch 2 | Batch 3 |
| --- | --- | --- |
| Rv_18S_rRNA_1 | Rv_YP_567084.1 | VvActinR |
| Rv_18S_rRNA_2 | Rv_YP_567084.2 | qVv-nad5-R |
| Rv_18S_rRNA_3 | Rv_YP_567072.1 | Rv_YP_567063.1 |
| Rv_18S_rRNA_4 | Rv_YP_567072.2 | Rv_XM_019225640.1 |
| Rv_18S_rRNA_5 | Rv_YP_567071.1 | Rv_YP_567056.1 |
| Rv_28S_rRNA_1 | Rv_YP_567103.1 | Rv_YP_567056.2 |
| Rv_28S_rRNA_2 | Rv_YP_567075.1 | Rv_YP_567106.1 |
| Rv_28S_rRNA_3 | Rv_YP_567107.1 | Rv_XM_019218790.1 |
| Rv_5S_rRNA_2 | RV_YP_002608363.1 | Rv_XM_002267690.4 |
| Rv_5_8S_rRNA_1 | Rv_YP_002608363.2 | Rv_XM_002284027.3 |
| Rv_5_8S_rRNA_2 | Rv_YP_567057.1 | Rv_YP_567061.1 |
| Rv_GQ849399.1 | Rv_XM_019216320.1 | Rv_YP_002608395.1 |
| Rv_KF544886.1 | Rv_YP_567108.1 | Rv_YP_567138.1 |
| 8S_rRNA_498_r | Rv_YP_567074.1 | Rv_YP_567083.1 |
| Rv_XM_002271755.3 | Rv_XR_002030730.1 | Rv_YP_002608358.1 |
| Rv_YP_567129.1 | Rv_XR_002031531.1 | Rv_YP_567064.1 |
| Rv_YP_002608376.1 | Rv_XR_002031185.1 | Rv_YP_567102.1 |
| Rv_NM_001281136.1 | Rv_XM_003632139.2 | Rv_XM_002285821.3 |
| Rv_YP_567131.1 | Rv_XM_019216227.1 | Rv_YP_567077.1 |
| Rv_XM_002278316.4 | Rv_YP_567133.1 | Rv_XR_004652793.1 |
| Rv_XM_019220594.1 | Rv_YP_567093.1 | Rv_XM_002267296.3 |
| Rv_XM_002271651.4 | Rv_YP_567059.1 | Rv_YP_002608358.1 |
| Ribo + genes | Chloro More expressed | Chloro More expressed 2 + actin + nad5 |

### 2.5. Real time quantitative polymerase chain reaction (qPCR)

All qPCR experiments were carried out in an ABI StepOne Plus Real Time PCR System (Applied Biosystems, California, USA). For each transcript quantification, three biological replicates were used in each treatment. qPCR data analysis and primer efficiencies were obtained using LinReg PCR software (Ruijter et al. 2009). Relative expression ratios and statistical analysis were performed using *R* software interface (Li et al., 2022), which uses the 2^−ΔΔCt^ method to analyze the relative changes in gene expression (Livak & Schmittgen 2001). A *p*-value below 0.05 was set as the cut-off for statistical significance.

### 2.6. Equation for Verification of Subtraction Efficiency

To ensure that the values obtained from our subtraction were objective, a modified ΔΔCt method was applied. The relative abundance of viral RNA was estimated based on the relationship between Ct values for the endogenous transcript (housekeeping) and the viral transcript, using the equation: 2^(Δ*Ctsws*(*eg*―*vg*)―Δ*Ctss*(*eg*―*vg*))^, where ‘eg’ corresponds to the endogenous transcript, ‘vg’ to the viral transcript, ‘sws’ to the sample without subtraction, and ‘ss’ to the subtracted sample (Figure 2c).

### 2.7. Sequencing and Bioinformatic Analysis RNA Sequencing

Total RNA was extracted from *Vitis vinifera* tissue samples as described in Section 2.2. The integrity and quality of the RNA were assessed by means of agarose gel electrophoresis and bioanalyzer (Agilent 2100) . The first four libraries were subtracted using RIBO Zero (Ribo-Zero^TM^ Magnetic Kit (Plant Leaf- Cat. No. MRZPL1224, Illumina, San Diego, CA, currently discontinued) and were subjected to Illumina Sequencing up to 100M reads per sample. The second set of four RNA libraries were prepared as follow: two controls—one substracted with the Ribo-Zero^TM^ as mentioned and the other without any depletion—and two experimental libraries using the Chloro-Zero methodology at “subtractor-to-subtracted” ratios of 1:1 and 2:1. Each library for this second round was sequenced on the Illumina platform to generate a minimum of 20 million paired-end reads, providing sufficient depth for comprehensive transcriptome analysis.

#### Quality Control and Preprocessing

Raw sequencing reads underwent quality assessment using FastQC (v0.11.9) to identify adapter contamination, low-quality bases, and overrepresented sequences. Subsequent trimming of adapters and low-quality bases was performed with Trimmomatic (v0.39) employing the following parameters: ILLUMINACLIP:TruSeq3-PE.fa:2:30:10, SLIDINGWINDOW:4:28, and MINLEN:50. Post-trimming quality was re-evaluated to confirm the effectiveness of the preprocessing steps.

#### Custom references Construction, Indexing, and Alignment

To enhance enrichment specificity, a series of custom references were constructed, incorporating highly expressed plastid, ribosomal, and nuclear genes, such as *psbA*, *rbcL*, ribosomal RNAs, as well as plant viral database. High-quality reads were aligned to the custom references and the *Vitis vinifera* genome (GenBank accession GCA_000003745.2) using Bowtie2 (v2.4.5) with sensitive mode parameters.

Alignment statistics, including mapping efficiency, concordant and discordant alignment rates, and coverage depth, were compiled using Samtools (v1.15.1). These metrics ensured the uniformity of alignments across libraries and allowed for an evaluation of the effectiveness of the Chloro-Zero subtraction protocol.

#### Transcriptomic Analysis

Reads from the RNA libraries of the first set of plant samples (CH40, MB21, 171, and P1103) were aligned to the *Vitis vinifera* reference genome (GenBank accession GCA_000003745.2) using Bowtie2 (v2.4.5) with sensitive mode parameters. Mapping statistics, including genome coverage and mapping efficiency, were calculated to evaluate transcript distribution and identify highly expressed genes. Transcript quantification was performed using eXpress (v1.5.1) (Roberts & Pachter, 2013) which enables rapid and accurate estimation of transcript abundances by modeling fragment assignment uncertainty.

In similar manners were treated the RNA reads from the MB21-derived subtraction treatments (second set of sequenced libraries, control without Ribo-Zero, Ribo-Zero-treated, and Chloro-Zero-treated at 1:1 and 2:1 subtractor-to-target ratios). The raw data were used for de-novo transcripts assembly by means of rnaviralSPAdes 3.15.3. From the assembled transcripts the viral ones were selected. The raw data of the four libraries (control without Ribo-Zero, Ribo-Zero-treated, and Chloro-Zero-treated at 1:1 and 2:1) were mapped against the same *Vitis vinifera* genome, the custom references transcripts and the reconstructed viral transcripts. This analysis enabled the assessment of subtraction efficiency by comparing transcript abundances, particularly for ribosomal RNAs, plastid genes, and viral RNAs.

## 3. RESULTS

### 3.1. Sensitivity Analysis and Probe Definition for Exome Depletion

To establish the detection limit of the NGS-based system, four test samples previously analyzed by gold-standard PCR were selected (Table 2). These samples, designated as CH40 (*Chardonnay*), MB21 (*Malbec*), and 171 (a recently described cultivar named *Criolla N°1*, Torres R et al., 2022), had been previously tested for the presence of *grapevine leafroll-associated virus 1-4* (GLRaV-1 to -4), *grapevine fanleaf virus* (GFLV), *grapevine fleck virus* (GFkV), *grapevine virus A* (GVA), *grapevine Pinot Gris virus* (GPGV), and *grapevine rupestris stem pitting-associated virus* (GRSPaV) (Table 2a). However, no prior information was available for sample P1103 (*Paulsen* 1103).

**Table 2.**
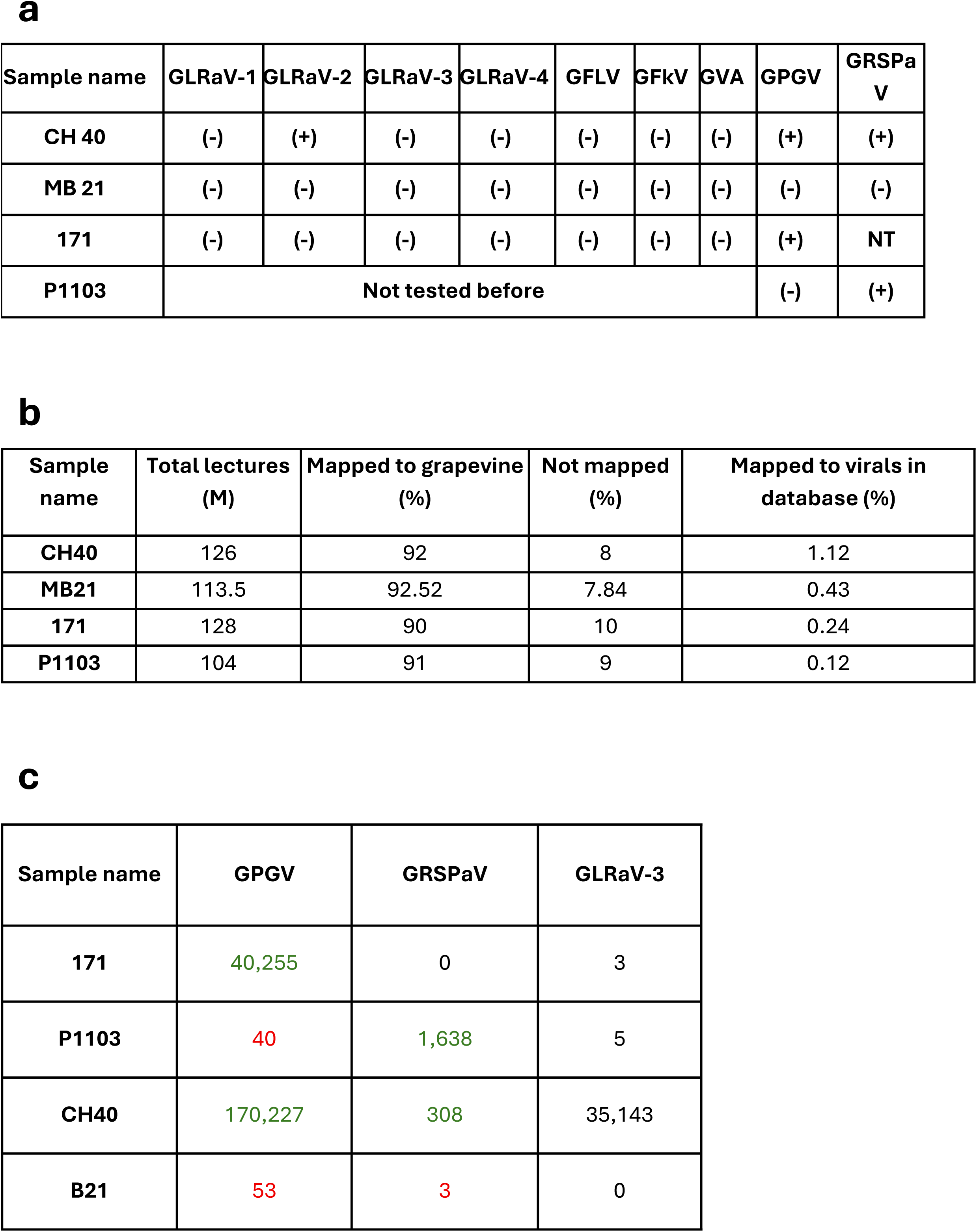
Analysis of samples by gold standard PCR and Next-Generation Sequencing (NGS). 2a. Grapevine plants whose viruses were characterized by standard PCR detection. These were GLRaV-1,2,3,4 (Grapevine Leafroll associated Virus), GFLV (Grapevine Fanleaf Virus), GFkV (Grapevine Fleck Virus), GVA (Grapevine Virus A). The only grapevine cultivar not tested before was the Paulsen P1103 rootstock. After sequencing, primers were designed to confirm the presence of GPGV (Grapevine Pinot gris Virus) and GRSaV (Grapevine Rupestris Stem pitting-associated Virus). **2b.** Reads exceeding 100 M per sample. Note that 90% or more of the mappings are associated with grapevine, 8% do not map to grapevine, and only 0.12% to 1.12% map to viral sequences. **2c.** Of the sequences that map to viruses, we could establish a detection limit more sensitive than a gold standard PCR. Viruses confirmed by PCR afterwards are marked in green, while those not confirmed are marked in red, establishing a new standard to define virus-free or virus-infected samples

| Sample name | GLRaV-1 | GLRaV-2 | GLRaV-3 | GLRaV-4 | GFLV | GFkV | GVA | GPGV | GRSPaV |
| --- | --- | --- | --- | --- | --- | --- | --- | --- | --- |
| CH 40 | (-) | (+) | (-) | (-) | (-) | (-) | (-) | (+) | (+) |
| MB 21 | (-) | (-) | (-) | (-) | (-) | (-) | (-) | (-) | (-) |
| 171 | (-) | (-) | (-) | (-) | (-) | (-) | (-) | (+) | NT |
| P1103 | Not tested before |  |  |  |  |  |  | (-) | (+) |

**b**
| Sample name | Total lectures (M) | Mapped to grapevine (%) | Not mapped (%) | Mapped to virals in database (%) |
| --- | --- | --- | --- | --- |
| CH40 | 126 | 92 | 8 | 1.12 |
| MB21 | 113.5 | 92.52 | 7.84 | 0.43 |
| 171 | 128 | 90 | 10 | 0.24 |
| P1103 | 104 | 91 | 9 | 0.12 |

**c**
| Sample name | GPGV | GRSPaV | GLRaV-3 |
| --- | --- | --- | --- |
| 171 | 40,255 | 0 | 3 |
| P1103 | 40 | 1,638 | 5 |
| CH40 | 170,227 | 308 | 35,143 |
| B21 | 53 | 3 | 0 |

All samples were sequenced with a depth exceeding 100 million reads per sample, ensuring robust genome coverage and reliable detection of viral sequences. As expected, approximately 90% of the reads aligned to the *Vitis vinifera* genome, while 10% remained unmapped. The remained reads were mapped to Plant Virus Database (http://47.90.94.155/PlantVirusBase/#/home) and between 0.1% and 1.1% were identified as viral or viroidal sequences (Table 2b).

Through comparative analyses, including *de novo* assemblies as previously described (e.g., Debat et al., 2019b), viral sequences were detected in all samples—even in those initially considered virus-free by PCR (Table 2c). For instance, Grapevine Pinot gris virus (GPGV) was identified in samples 171 and CH40, with over 40,000 reads mapped to this virus, consistent with PCR results. Similarly, low levels of *Grapevine Rupestris stem pitting-associated virus* (GRSPaV) were detected, ranging from 300 to 1,600 reads in CH40 and P1103 (Table 2c). On the other hand, low amounts of GPGV sequences were detected in samples P1103 and MB21 (40 and 53 reads, respectively), despite the absence of amplification in the corresponding PCR tests. A similar case was observed for GRSPaV in sample MB21, where only 3 reads were detected (Table 2c).

To confirm the presence of the detected viruses, specific primers were designed for PCR validation. Viral sequences detected by NGS and confirmed by PCR are indicated in green, while those undetected by PCR are marked in red (Table 2c). The case of sample CH40 and GLRaV-3 appears to be different. Although the virus was not detected by PCR but detected by NGS, the high number of reads mapped to GLRaV-3 suggests that this is more likely a PCR false negative error rather than a sensitivity issue. These results established two key outcomes, the identification of a candidate plant sample with minimal viral content to be used as a pure grapevine transcriptome (MB21), and the estimation of the sensitivity threshold of the NGS technique, which, under the experimental conditions applied, outperformed conventional gold-standard PCR methods in terms of accuracy.

### 3.2. First Methodology for Subtraction

Using RNA extracted from the MB21 sample—characterized by a minimal viral load—we implemented the subtraction protocol described in Figure 1. The method involved synthesizing biotin-dUTP-labeled probes from the complete grapevine transcriptome, hybridizing them to the RNA targets, and subsequently removing the resulting RNA–probe hybrids with streptavidin-coated magnetic beads. All steps were conducted successfully and without technical issues, confirming the feasibility of the approach.

The results of the genomic RNA subtraction, summarized in Table 3, reveal variability in subtraction efficiency across different experimental rounds. Efficiency was estimated by comparing Ct values for endogenous reference genes and viral targets between subtracted and non-subtracted RNA samples (see formula in the Materials and Methods section). In the first round, a ΔCt of 1.18 was observed in the T sample (Torrontés, high viral load), when using ST351 probes at a subtractor-to-sample ratio of 3.5:1—corresponding to a 2.26-fold enrichment of viral RNA. Similar results were obtained in round four (ΔCt = 1.32; 2.49-fold enrichment). In rounds eight and ten, subtraction efficiency improved further, with ΔCt values of 1.93 and 1.92, equivalent to 3.81- and 3.78-fold enrichment, respectively.

**Table 3.**
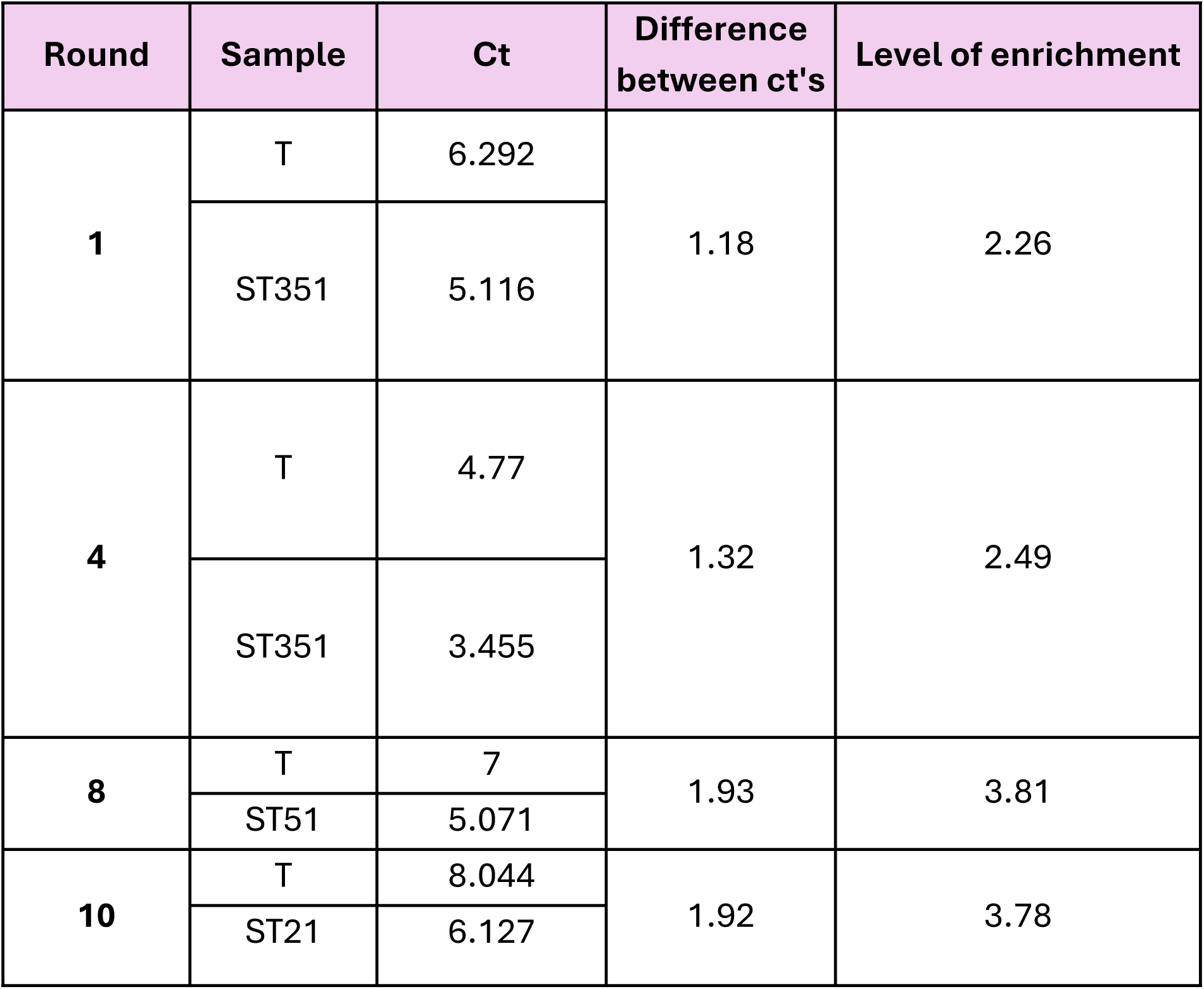
Results of subtractions with random primers and Oligo (dt). Differences in Ct values between control T (Torrontés) and the corresponding subtraction were observed in subtraction rounds 1, 4, 8, and 10: in round 1, the subtraction ratio was 3.5:1 between subtractor and subtracted, the same for round 4. In round 8, the ratio was 5:1, and in round 10, it was 2:1.

These findings demonstrate that the protocol enables enrichment of viral RNA to a degree that could allow the sequencing of up to four samples per run—compared to the single-sample limitation of conventional methods—while maintaining similar sensitivity and significantly reducing costs.

However, the variability observed across experimental replicates highlighted limitations in the reproducibility of the method (Table 3 shows only 4 out of 10 assays). While the results represent a step forward in achieving efficient and cost-effective viral exome enrichment, further optimization is required to improve both the reliability and consistency of the subtraction process, ensuring more effective and reproducible concentration of viral RNA while minimizing host RNA contamination.

### 3.3. Characterization of Transcripts for Improving Subtraction Probes

To refine subtraction efficiency, transcriptomic data from all four sequenced samples (Table 2) were further analyzed to identify the highly expressed genes. The analysis revealed that all samples exhibited similar expression profiles, with *psbA* (Photosystem II protein D1) being the most abundant transcript (Table 1a). It should be noted that all sequencing libraries analyzed in this section were subjected to rRNA depletion using the Ribo-Zero method prior to sequencing. This extremely high expression level, which is consistent with previous findings by Forsythe et al. (2022), underscores the dominance of plastid transcripts in the cellular mRNA pool. Other highly expressed plastid genes, such as *rbcL* (RuBisCO large subunit), were also observed, alongside ribosomal RNA (rRNA), whose abundance further highlights the challenge of efficient host RNA depletion. The full list of the 200 most expressed genes is provided in Supplementary Table 2.

Building on this analysis, three probes batches were designed (Table 1b): Batch 1, containing 13 rRNA genes and other 10 highly expressed genes from the top 200 list; Batch 2, comprising highly expressed plastid and mitochondrial genes identified in the transcriptomic analysis; and Batch 3, consisting endogenous highly expressed genes along with endogenous nuclear housekeeping genes such as *actin* and *nad5*. Primer batches were designed based on the selected genes and subsequently used to synthesize cDNA probes targeting the most abundant transcripts. This approach was aimed to increased subtraction efficiency and addressed the limitations inherent to probe synthesis strategies relying on random primers or oligo(dT).

To validate the effectiveness of the improved probes set in the subtraction efficiency, specific PCR primers were designed for key targets genes in each batch (Supplementary Table 1). Genes highlighted in bold were used to assess transcript depletion and to evaluate probe performance. The complete list of primers for probes cDNAs synthesis is provided in Supplementary Table 1B, offering a comprehensive resource for future applications of this methodology.

### 3.4. Exome Rejection with Specific Primers (Chloro-Zero)

The three primers batches were mixed in a 1:1:1 ratio to generate biotin-labeled cDNA probes. These probes, collectively referred to as “Chloro-Zero,” were used in the hybridization and pull-down steps outlined in Figure 2a and evaluated for their efficiency in depleting highly abundant transcripts. The probes were applied at two subtractor-to-target RNA ratios, 1:1 and 2:1, to assess the subtraction efficiency and reproducibility. This design aims to optimize RNA subtraction without unintentionally depleting viral RNA.

The subtraction process was quantified by RT-qPCR, comparing the abundance of endogenous transcripts (*actin*, *nad5*, *rbcL*, *28s*, and *psbA*) against a region of *GLRaV-2* genome. Figure 2b presents qPCR results for the Actin housekeeping gene and a viral gene, comparing RNA samples before and after subtraction. The data reveals a more pronounced depletion of Actin transcripts relative to the viral gene. The full results, summarized in Figure 2c, revealed that enrichment efficiency varied depending on the endogenous gene and subtraction ratio. At the 1:1 ratio, psbA and 28s showed significant reductions of 2.07-fold and 274.72-fold, respectively. At the 2:1 ratio, the same genes showed reductions of 2.52-fold and 132.20-fold, respectively. Other genes, such as *actin7* and *nad5*, exhibited lower but consistent reductions across both ratios, ranging from 1- to 5-fold. Notably, 28s showed the highest subtraction efficiency, with reductions exceeding 270-fold under the 1:1 ratio.

The formula used to calculate enrichment efficiency (Figure 2c) ensured accuracy by accounting for the ΔCt differences between endogenous and viral genes under subtraction or non-subtraction conditions. This approach minimized the risk of artifacts that could result from subtractor probes unintentionally binding to viral transcripts. The results consistently demonstrated significant depletion of abundant transcripts while preserving viral RNA across all replicates, confirming the robustness and reproducibility of the method.

Overall, the Chloro-Zero system effectively reduced host RNAs, enabling the enrichment of viral RNA for downstream analyses. These positive results pave the way for the next phase of research, focusing on RNA integrity and virome profiling using next-generation sequencing (NGS).

To complement the qPCR-based validation of the exome subtraction system and provide an additional layer of analysis, capillary electrophoresis using the Bioanalyzer system was incorporated. The evaluation included a non-subtracted control, and samples subjected to 1:1 and 2:1 subtraction ratios. Bioanlayzer electropherograms revealed a marked reduction in ribosomal RNA peaks (18S and 25S) in the subtracted samples, confirming the effective removal of rRNA by the Chloro-Zero system (Figures 3b and c). As expected, the RNA Integrity Number (RIN) was only calculated for the intact control sample, where the presence of well-defined rRNA peaks allowed for a reliable assessment of RNA integrity (Figure 3a). Furthermore, the control sample enabled a direct comparison between the Chloro-Zero method and the commercial Ribo-Zero Plus rRNA Depletion Kit (Illumina©). Results suggest that the Chloro-Zero system constitutes a cost-effective alternative for reducing rRNA, tailored to the specific experimental objective of minimizing sequencing costs without compromising RNA quality.

**Figure 3.**
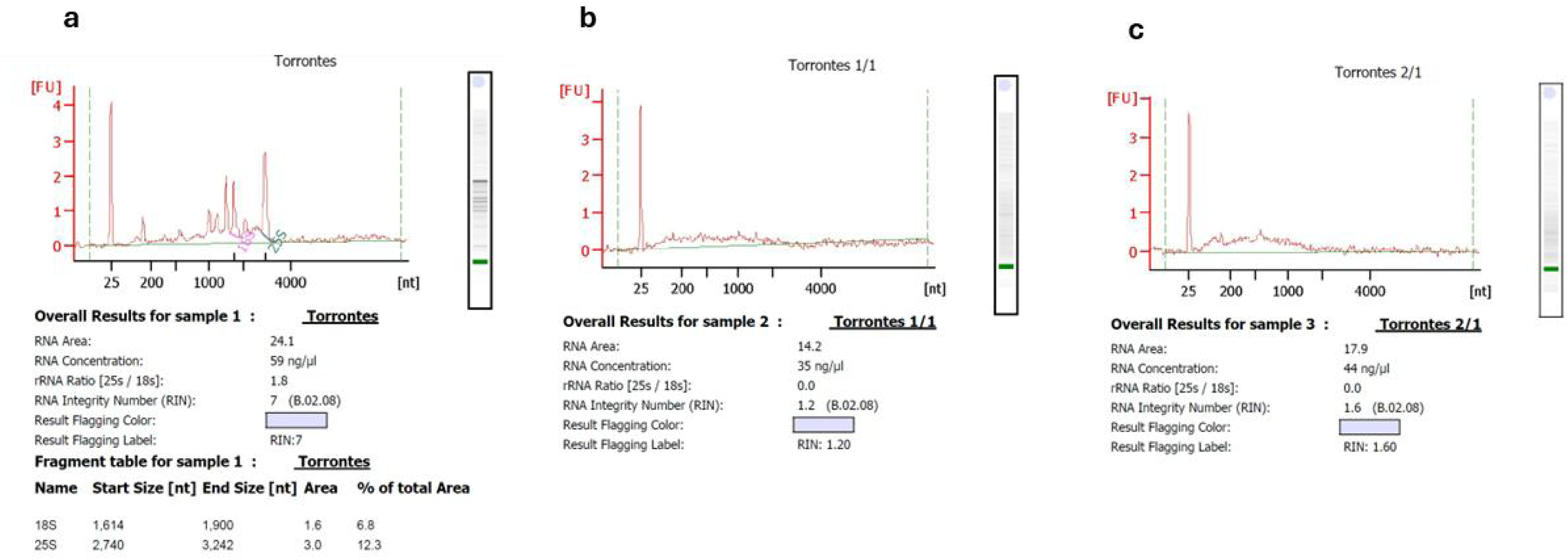
Contrast between control and subtracted (enriched) samples. Electrophoresis analysis (Bioanalyzer) of RNAs from **a.** Sample of the Torrontés cultivar. **b** and **c**, same sample analyzed after performing exome rejection in a subtraction ratio of 1:1 and 2:1 between subtractor (biotinylated designed probes) and subtracted (target RNA). Note that in both graphs **b** and **c**, unlike in **a**, the peaks associated with ribosomal 18s and 25s disappear, suggesting that these have been eliminated or degraded.

### 3.6. NGS Analyzed Samples and Enrichment Efficiency

To evaluate the performance of the different tested RNA subtraction strategies, a RNAseq assay was set, four libraries using the same RNA sample were sequenced: a negative control (without Ribo-Zero), a commercial control (with Ribo-Zero subtractions), and two experimental treatments using Chloro-Zero at subtractor-to-target ratios of 1:1 and 2:1. Table 4a summarizes the RNA-seq results, including total read counts, mapping rates to the Vitis vinifera chloroplast and mitochondrial genomes, rRNAs genes and remaining Vitis nuclear genome. Comparison of Chloro-Zero–treated samples (ratios 1:1 and 2:1) with the negative control revealed a measurable effect on transcript depletion. In particular, the 2:1 Chloro-Zero treatment resulted in approximately a 10% reduction in chloroplast and in mitochondrial genome content (Table 4a). This treatment also contributed to a modest reduction in non-coding and highly expressed rRNAs. Nevertheless, Ribo-Zero® exhibited a stronger overall depletion efficiency.

**Table 4.**
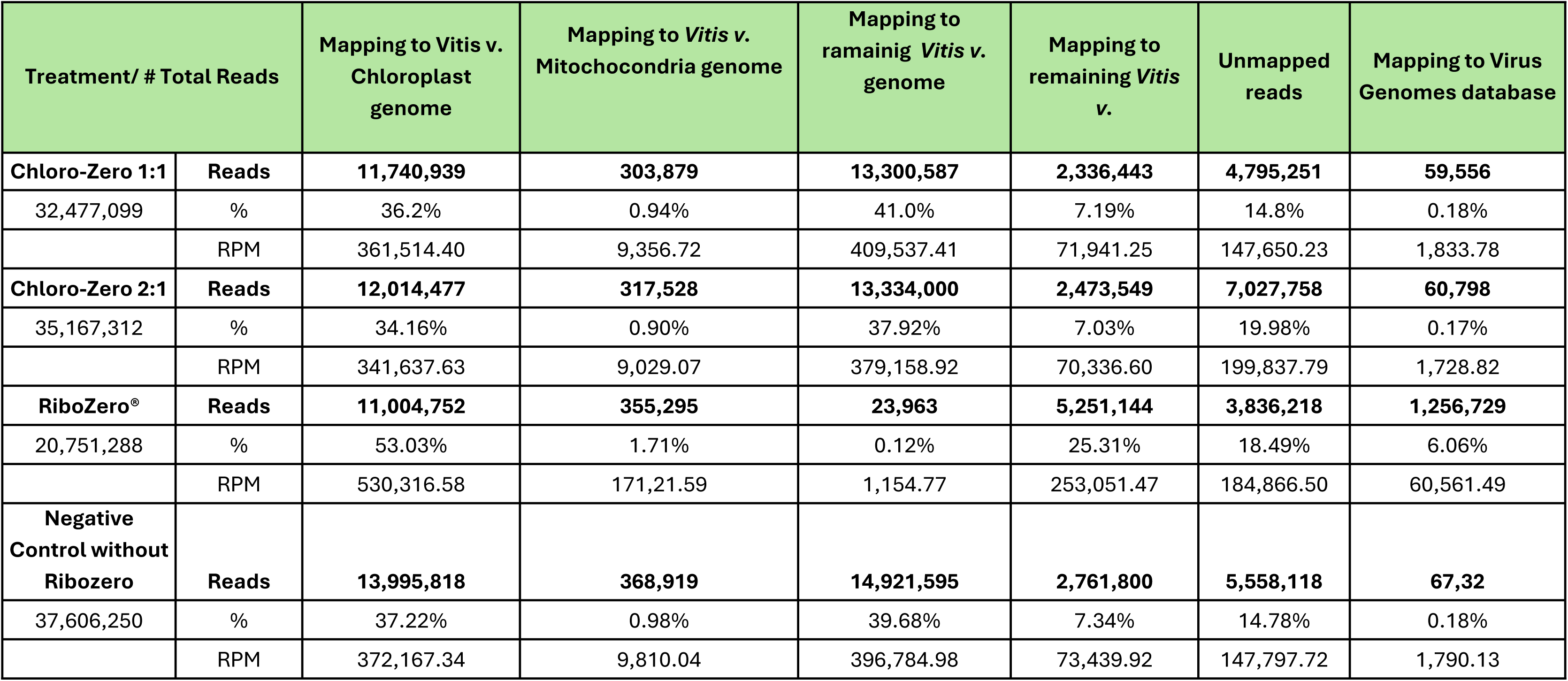
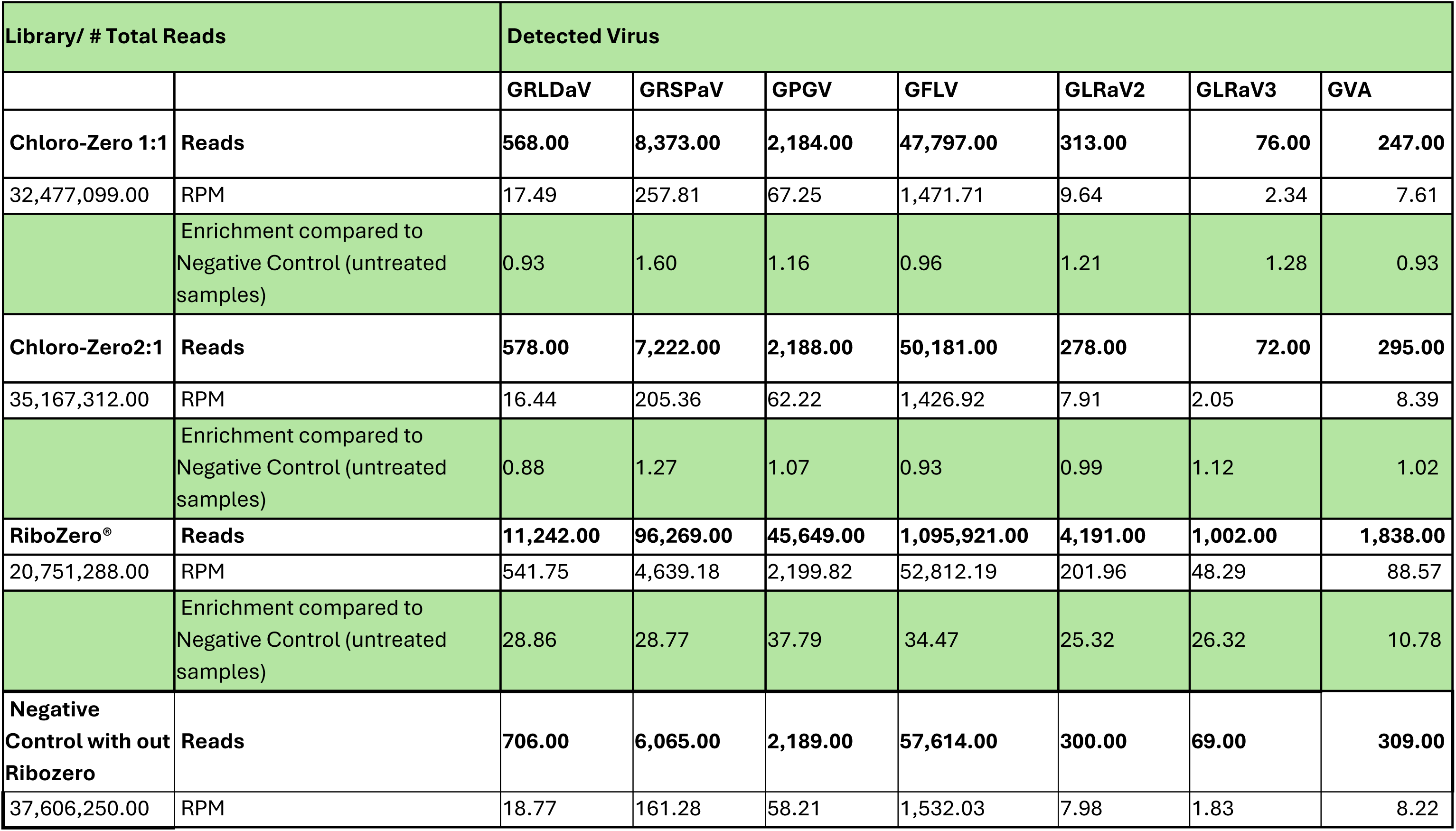
Overview analysis of samples analyzed by NGS. This table presents sequencing results obtained from the same RNA sample under four conditions designed to evaluate different RNA subtraction strategies: a negative control without Ribo-Zero, a commercial control with Ribo-Zero subtraction, and two experimental treatments using Chloro-Zero at subtractor-to-target ratios of 1:1 and 2:1. Panel (a) shows general mapping statistics for the sequenced samples. Panel (b) compares viral genome detection across seven grapevine viruses (GRLDaV, GRSPaV, GPGV, GFLV, GLRaV2, GLRaV3, and GVA) among the four conditions.

Furthermore we were able to detect 7 viral RNAs in the test samples, resulting in the identification of grapevine Roditis leaf deformation virus, grapevine rupestris stem pitting virus, grapevine Pinot Gris virus, grapevine fanleaf virus, grapevine virus A, grapevine satellite virus, grapevine leafroll associated virus 2 and 4. Surprisingly not all of them were enriched in a similar way. Chloro-Zero treatments showed their potential by enriching some viral genomes RNA approximately up to 60% at the better ratio showed to GRSPaV. (Table 4b)

While not as effective in depleting highly abundant rRNAs as Ribo-Zero (note, this commercial product is no longer available as standalone reagent), the Chloro-Zero system preserved and enhanced viral RNA representation, a key feature for virome profiling. These results shows Chloro-Zero idea as a promising, cost-effective alternative for enriching low-abundance transcripts, particularly viral RNA, without requiring extensive depletion of all highly expressed RNAs.

### 3.7. Analysis of the Increase in Subtraction Ratio

To evaluate whether the efficiency of subtraction could be enhanced by modifying the ratio between subtractor and target RNA, we conducted a final test using a substantially higher subtractor-to-target ratio of 10:1. Endogenous reference genes, including actin and several 18S rRNAs, were used as internal benchmarks and compared to GLRaV-2 gene to assess the effectiveness of the subtraction. Supplementary Table 3, exhibited a reduction in expression of ribosomal RNA genes (e.g., 18S-2 and 28S) by 3 to 4-fold under these conditions. However, these results lacked reproducibility, with inconsistent outcomes observed even at the same 10:1 ratio, and further attempts at a 50:1 ratio (Supplementary Table 3) did not improve reliability.

These findings indicate that increasing the concentration of the subtractor relative to the target viral RNA does not enhance the efficiency of the Chloro-Zero system.

On the contrary, such adjustments can negatively impact the methodology by introducing variability and escalating costs due to the higher consumption of labeled probes and streptavidin reagents.

Nevertheless, the ChloroZero system remains a promising approach for transcript subtraction and enrichment. Its ability to target and slightly enrich viral RNA, as demonstrated in earlier tests, highlights its potential for applications requiring cost-effective and customizable methodologies. The flexibility of Chloro-Zero, coupled with its scalability, positions it as an adaptable solution for transcriptomic studies focused on enriching low-abundance RNAs, particularly in resource-limited scenarios. Future refinements in probe design and application protocols may address the reproducibility challenges observed at high subtraction ratios, further unlocking the system’s potential for virome profiling and other RNA enrichment applications

## 4. DISCUSSION

Next Generation Sequencing (NGS) has become an essential tool for the detection and characterization of plant viruses (Navrotskaya et al., 2021; Eichmeier et al., 2016), particularly in crops like *Vitis vinifera*, where mixed infections and asymptomatic viral reservoirs pose significant challenges for disease management. Despite its advantages, a major limitation of RNA sequencing in virome studies is the overwhelming presence of highly expressed host transcripts—particularly non-coding RNAs such as ribosomal RNAs and the unexpectedly abundant plastid-derived mRNAs (Forsythe et al., 2022). These transcripts dominate sequencing reads, often masking low-abundance viral sequences. Commercial rRNA depletion kits, like Ribo-Zero Plant, have been discontinued by the original manufacturer and are no longer available as standalone reagents (<u>supportassets.illumina.com</u>). Additionally, the lack of plant-specific formulations in current alternatives underscores the need for developing tailored strategies for plant transcriptomes. To address these challenges, this study explored the development of a targeted subtraction approach, leveraging transcriptomic data to design customized probes capable of selectively depleting highly expressed host RNAs while preserving viral RNA.

The candidate genes selected for probe design were derived from RNA-seq data of *V.vinifera* samples, ensuring an evidence-driven approach. The most highly expressed transcripts, including *psbA* (Photosystem II protein D1), *rbcS* (RuBisCO small subunit), and multiple rRNA sequences, were consistently among the most abundant across independent biological replicates. These findings were corroborated by previous studies (Forsythe et al., 2022), reinforcing the reliability of transcript selection. Given that high transcript abundance correlates with high sequencing representation, novel transcriptomic analyses in grapevine are expected to yield similar rankings, with only minor variability on relative expression levels.

While the probe-based depletion approach holds theoretical promise, the observed rRNA depletion was limited under the conditions tested. The absence of rRNA peaks in electrophoresis profiles and chromatograms initially suggested successful subtraction, yet NGS data revealed persistent rRNA presence, with mapping percentages exceeding 50% in all Chloro-Zero-treated samples. This discrepancy aligns with previous reports showing that rRNA contamination can remain undetectable in electrophoresis yet still be present in sequencing reads (Zhao et al., 2018). These findings highlight the importance of complementary validation methods beyond gel electrophoresis when assessing depletion efficiency.

To optimize the protocol, multiple parameters were systematically varied, including subtractor-to-target RNA ratios, probe concentrations, and streptavidin bead volumes. The experimental design ensured precise control over these variables, with rigorous adherence to hybridization and pull-down conditions. However, even at increased subtractor ratios of 10:1 and 50:1, no consistent improvement in rRNA depletion was observed. These results suggest that the hybridization efficiency or probe specificity may be limiting factors, rather than subtractor concentration alone.

The limited efficiency of rRNA depletion observed through NGS suggests that electrophoresis data may provide misleading results. A plausible explanation is that rRNA molecules were partially degraded—producing no visible peaks on electrophoresis gels yet containing fragments that remained detectable during sequencing and contributed to the residual rRNA signal. This latter hypothesis also correlates with qPCR analysis.

Despite these challenges, one of the key findings of this study was the ability of the ChloroZero protocol to enrich viral RNA. While depletion of host transcripts was limited, viral reads, particularly those corresponding to *GLRaV-2, -4 and GRSPaV*, were fairly enriched in ChloroZero-treated samples. Given that RiboZero® is optimized for animal and bacterial rRNA depletion and lacks plant-specific formulations, its efficiency in plant transcriptomes remains suboptimal. The development of organism-specific depletion strategies, such as the one tested here, remains a promising direction for future research.

To further refine this methodology, improvements in probe synthesis and hybridization conditions could be explored. Outsourcing the production of biotin-labeled probes to specialized synthesis services could enhance quality control, ensuring consistent probe-to-target hybridization. Future protocol modifications could include replacing streptavidin bead-based pull-down with enzymatic degradation of hybridized fragments. Alternatives such as DNase II treatment may offer selective RNA degradation while preserving viral RNA integrity, partly due to the structured nature of viral genomes (Zhuang et al., 2024).

In summary, this study highlights the complexities of transcript depletion in plant virome studies and underscores the challenges associated with developing cost-effective alternatives to commercial depletion kits. While the tested approach did not achieve significant rRNA removal, it demonstrated the ability to enrich viral RNA, making it a valuable framework for further optimization. Future refinements in probe design, hybridization conditions, and bioinformatics analyses may ultimately lead to a more effective and reproducible subtraction method, tailored specifically for plant transcriptomes.

## Acknowledgements

This work was supported by Instituto Nacional de Tecnología Agropecuaria (INTA) Grant# PD-INTA: 2023-PD-L01-I083, Agencia Nacional de Promoción de la Investigación, el Desarrollo Tecnológico y la Innovación Grants # PID 2018 -021. Additional funding was provided by Catena Institute of Wine.

---

**Supplementary Table 1.**
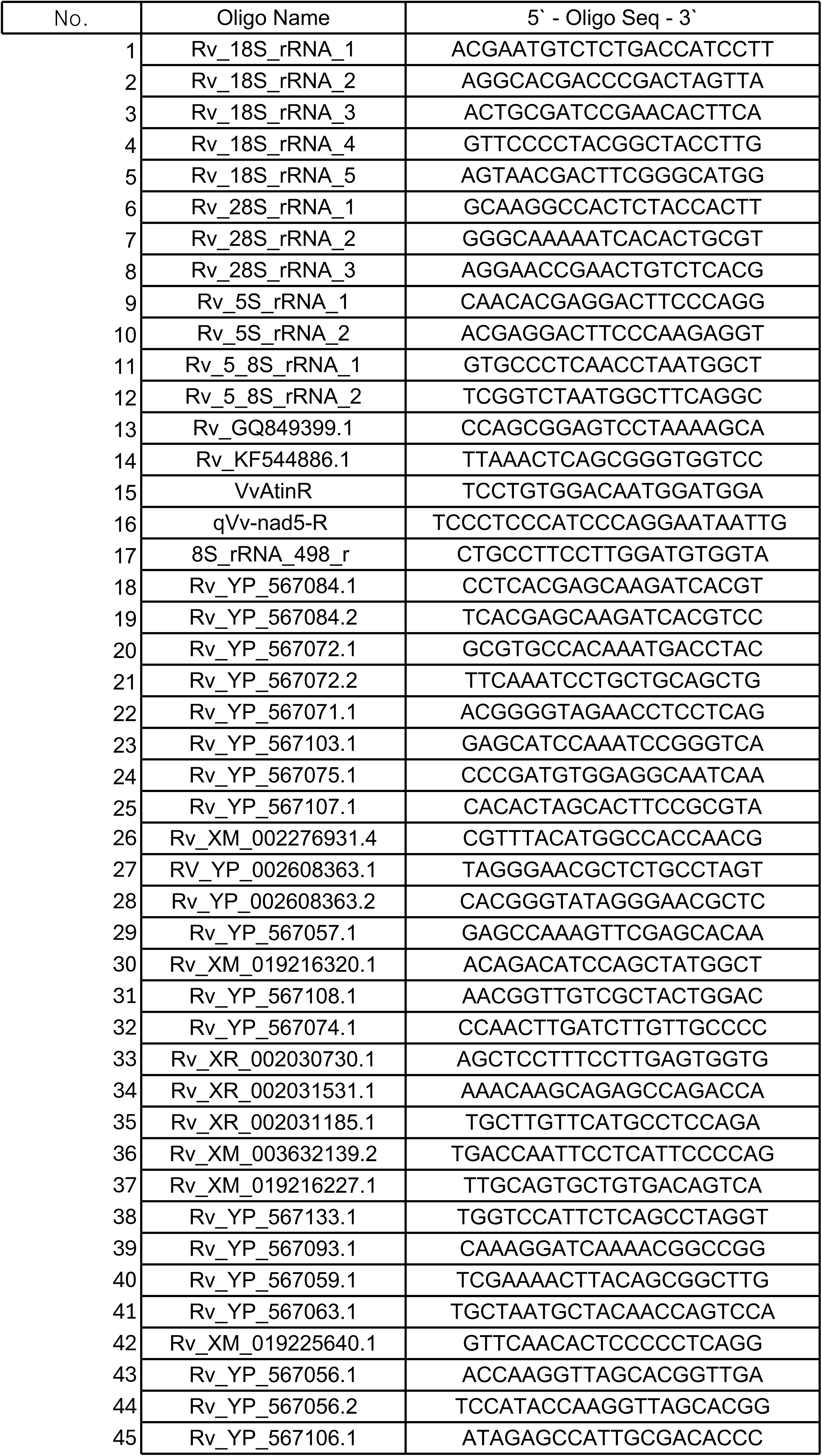
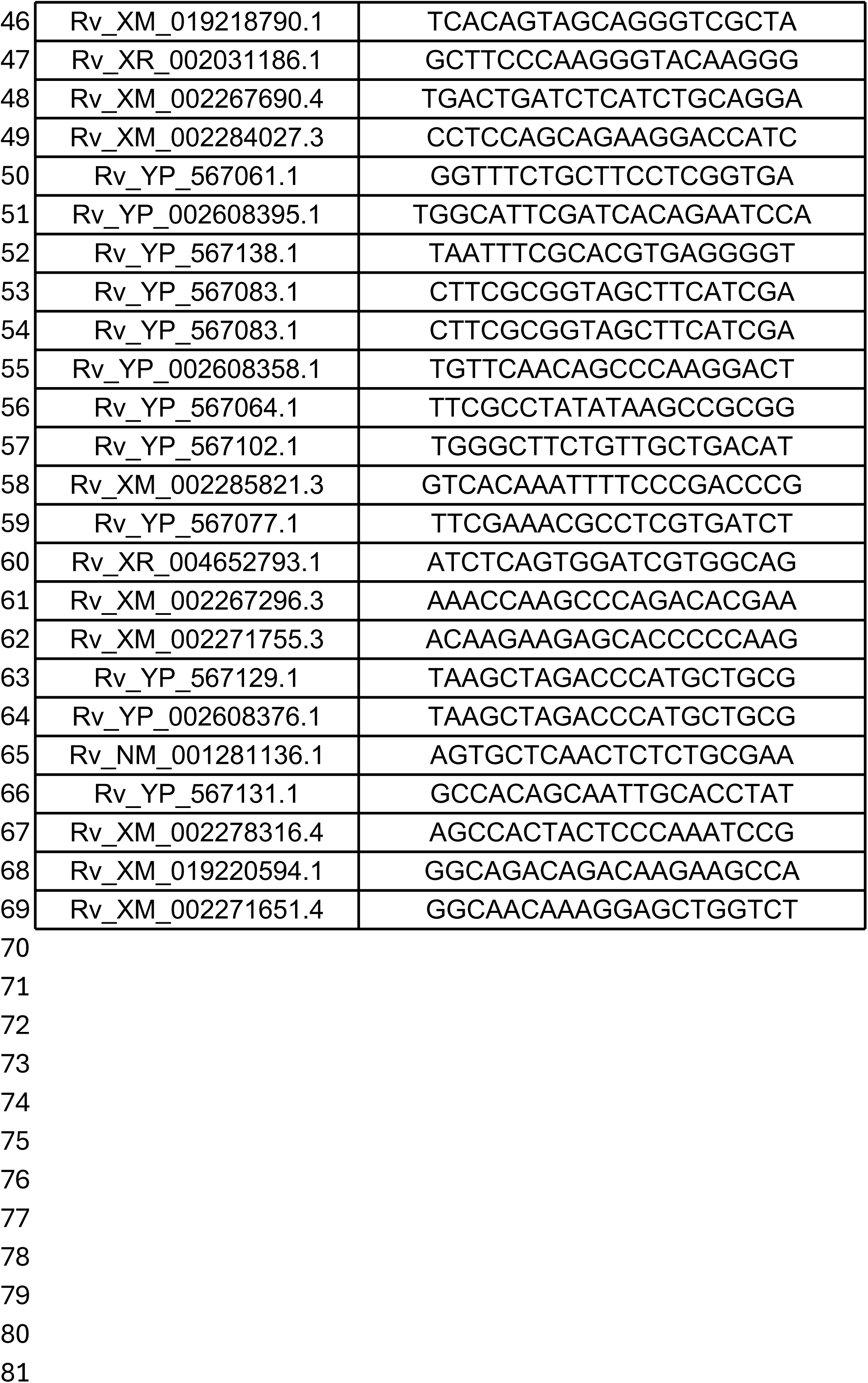
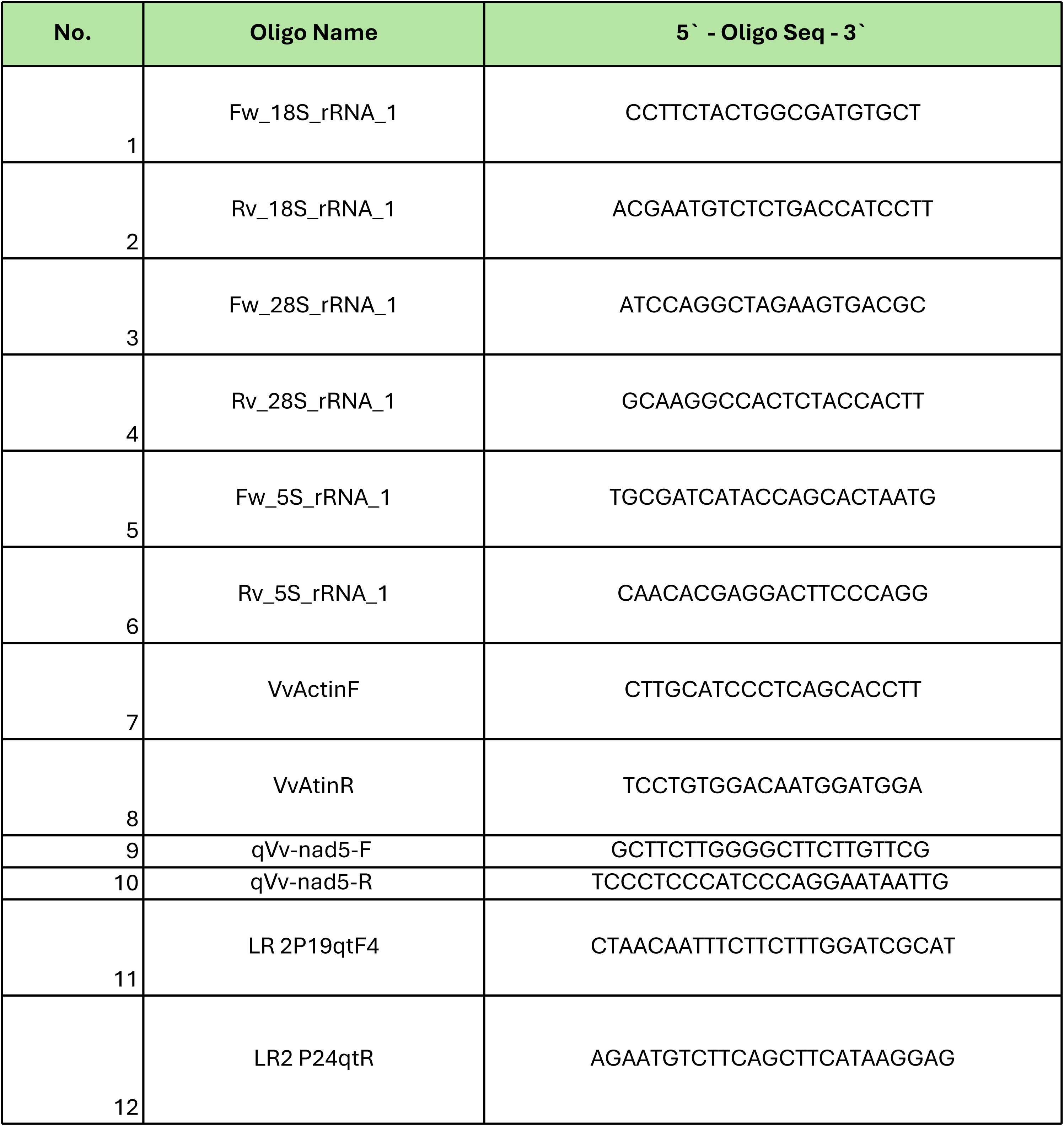
Primers sequence list, A Chloro-Zero primers. B qPCR primers.

**Supplementary Table 2.**
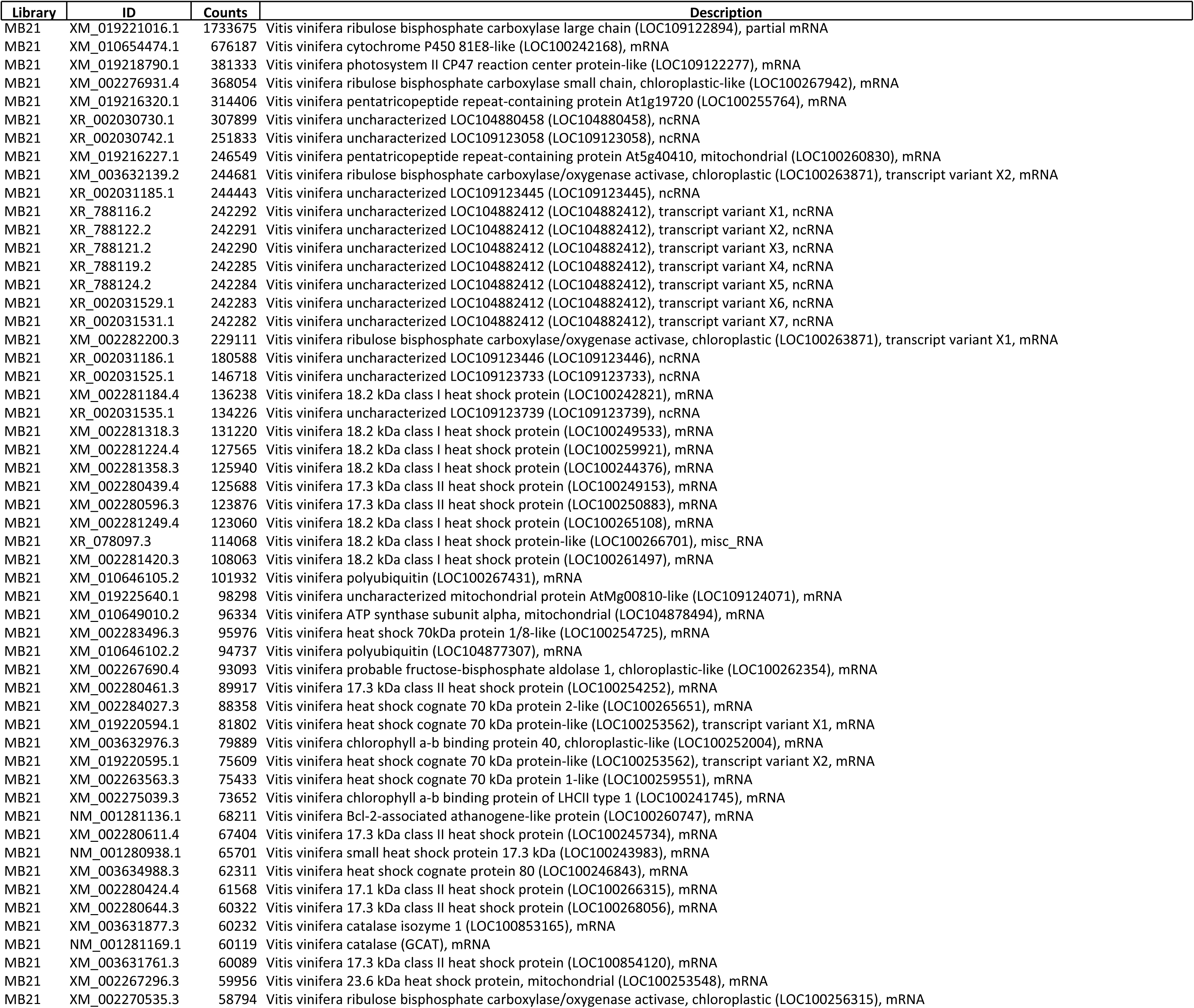
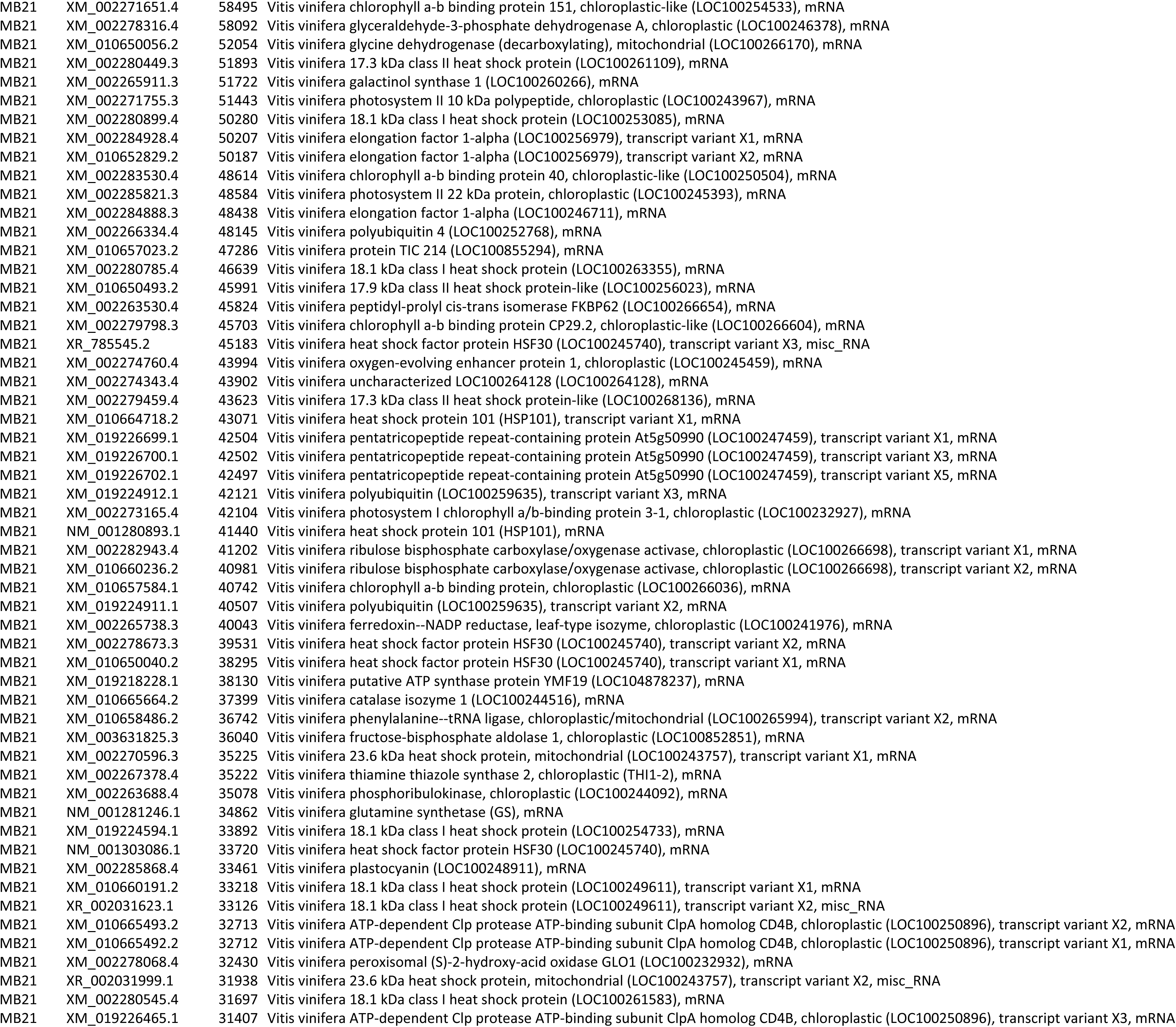
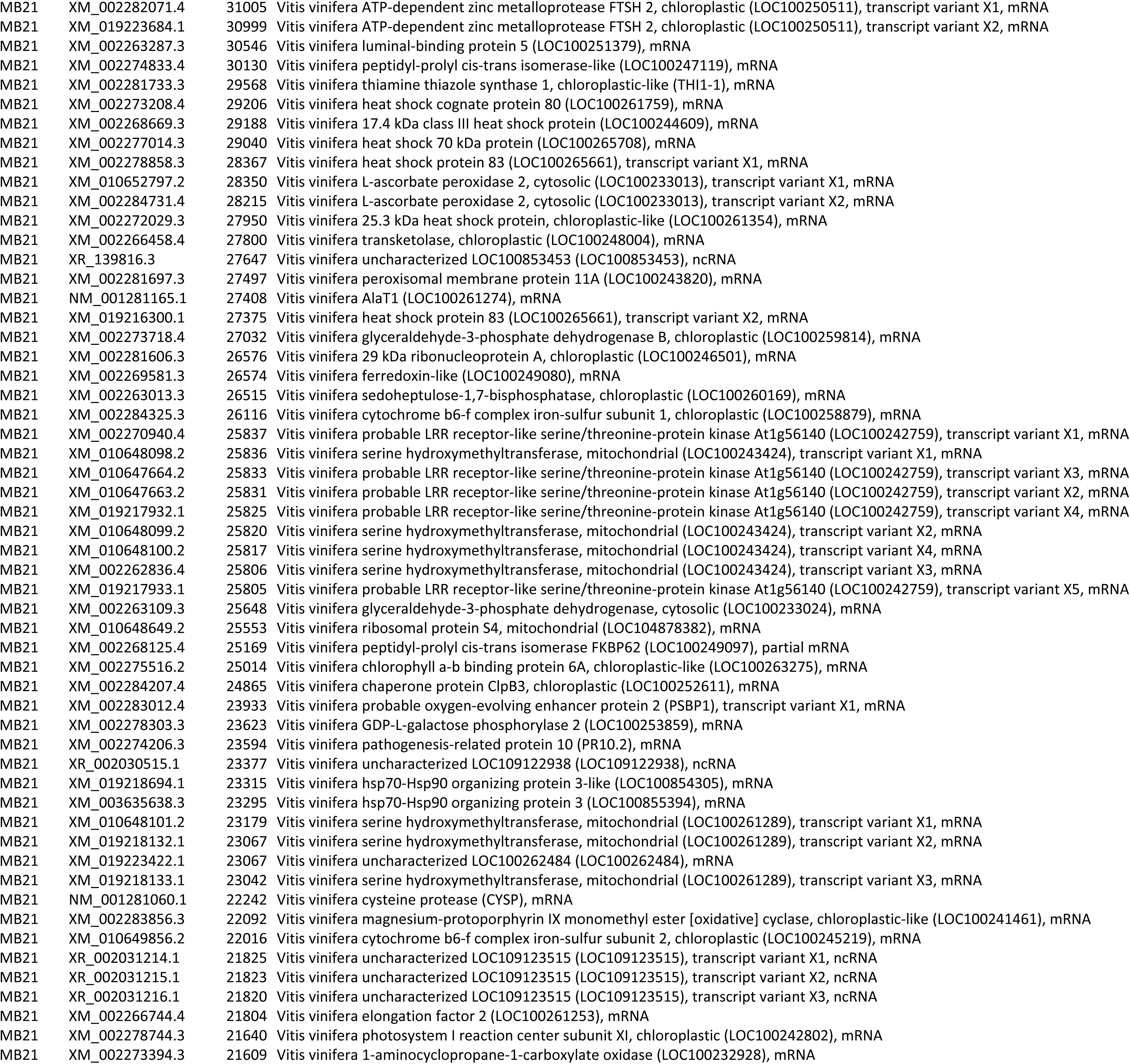
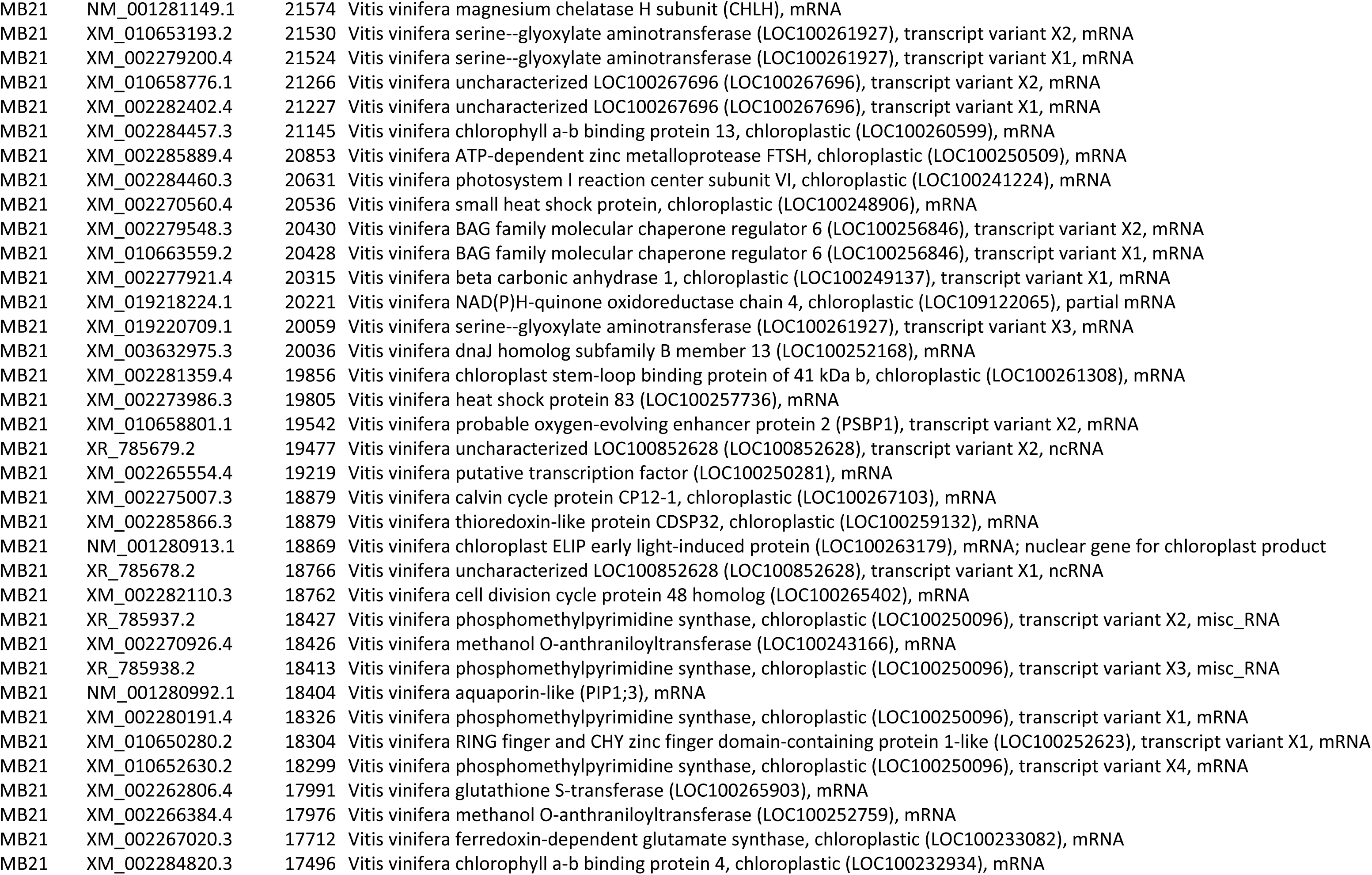
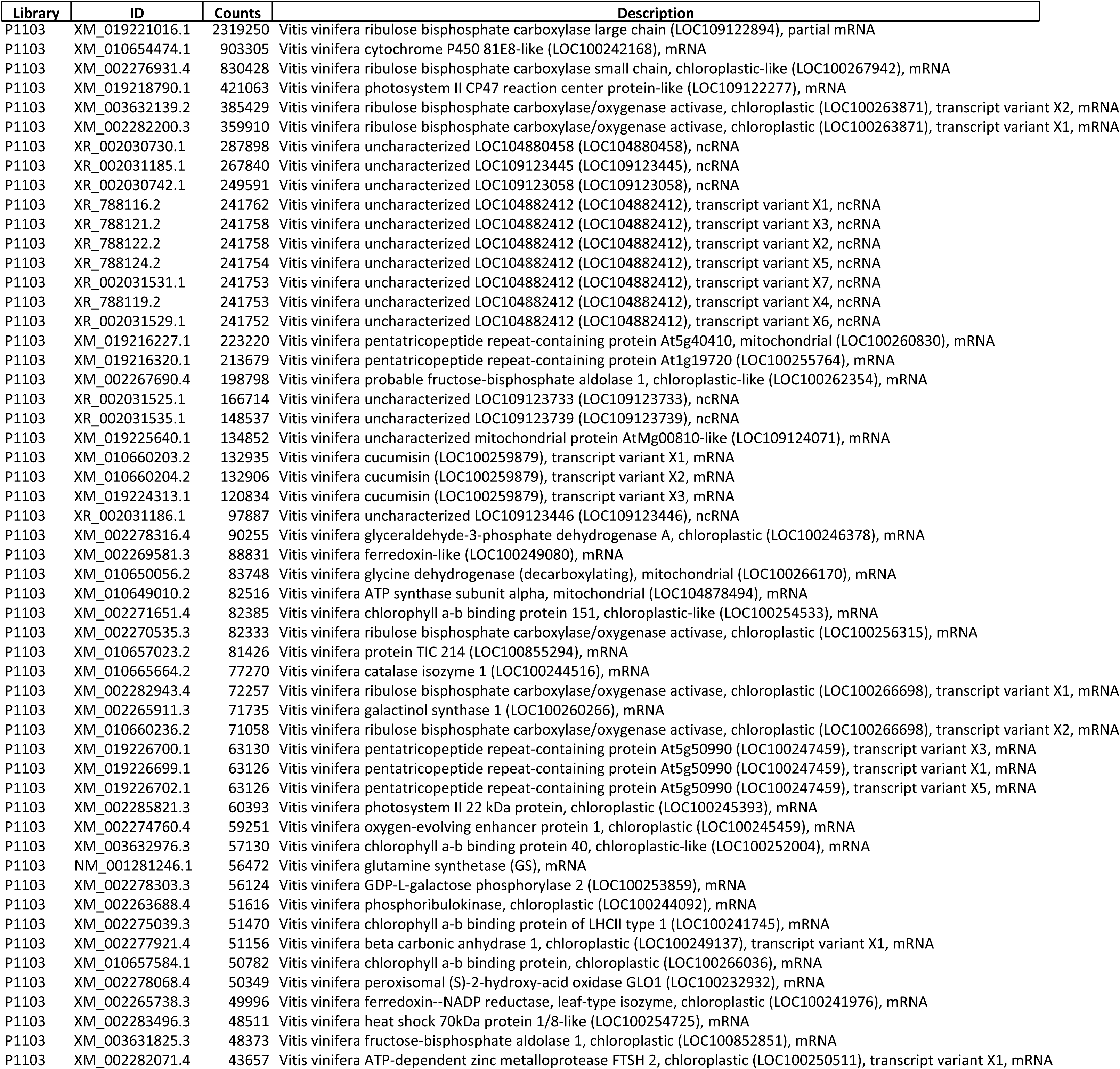
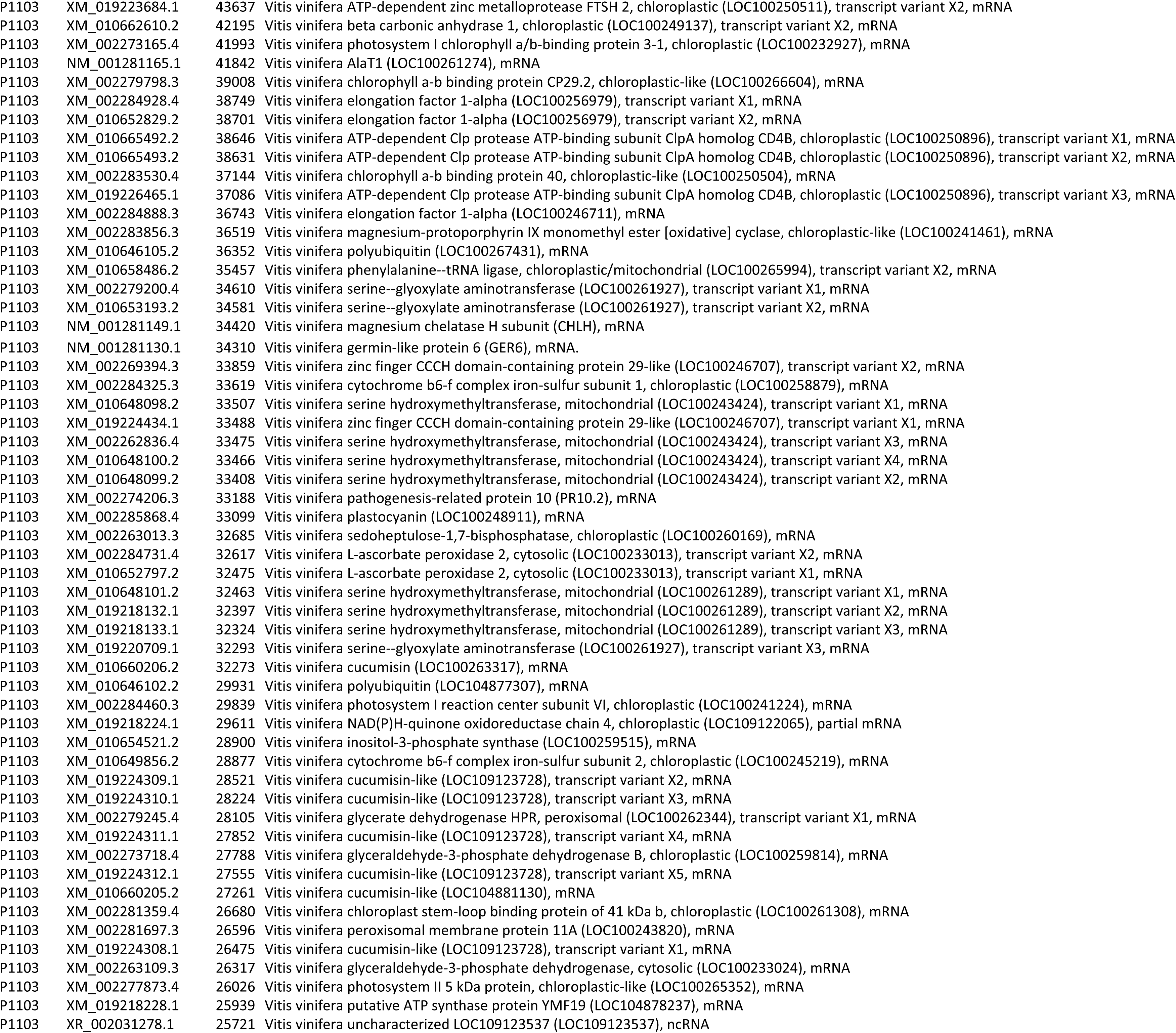
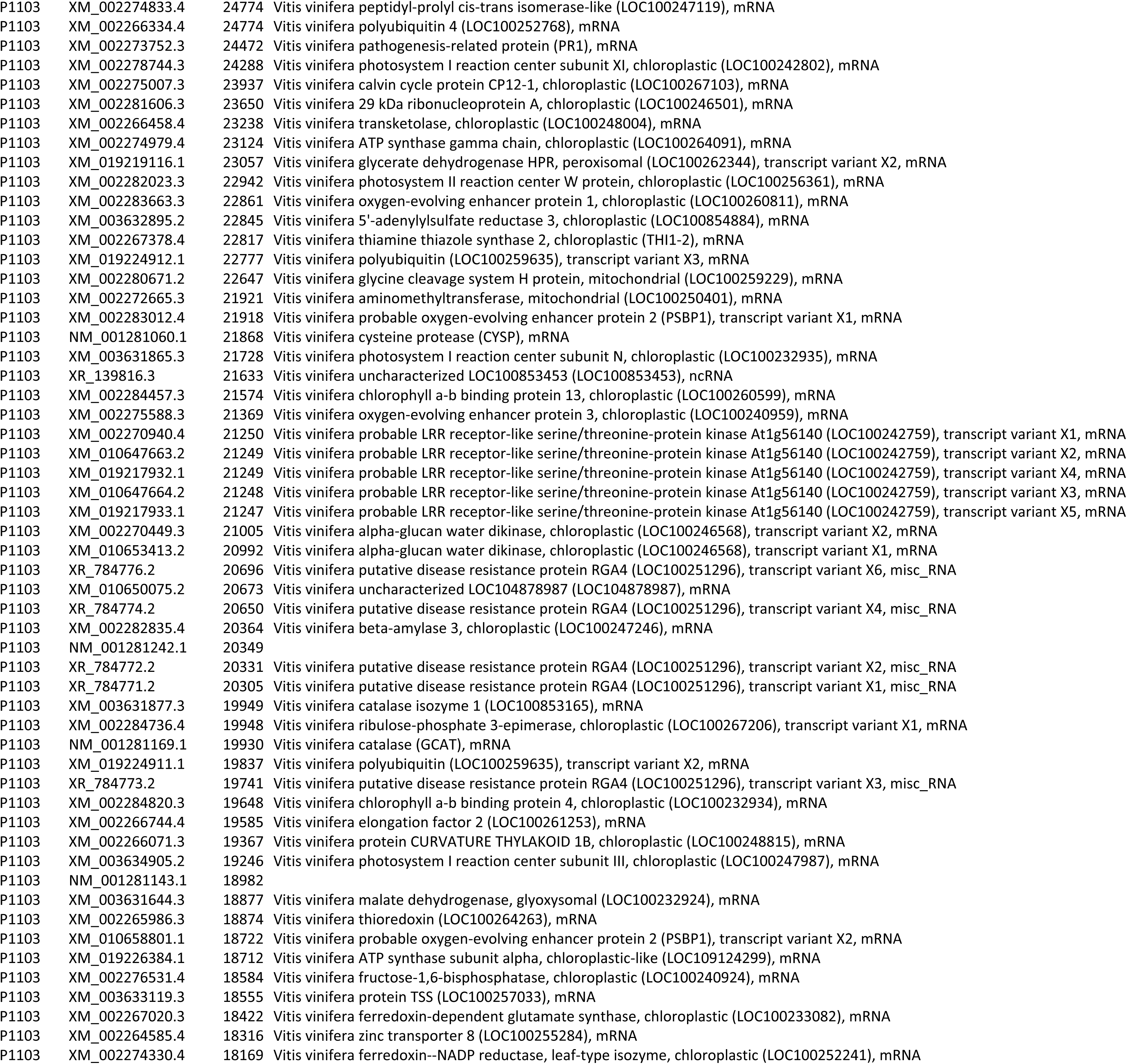
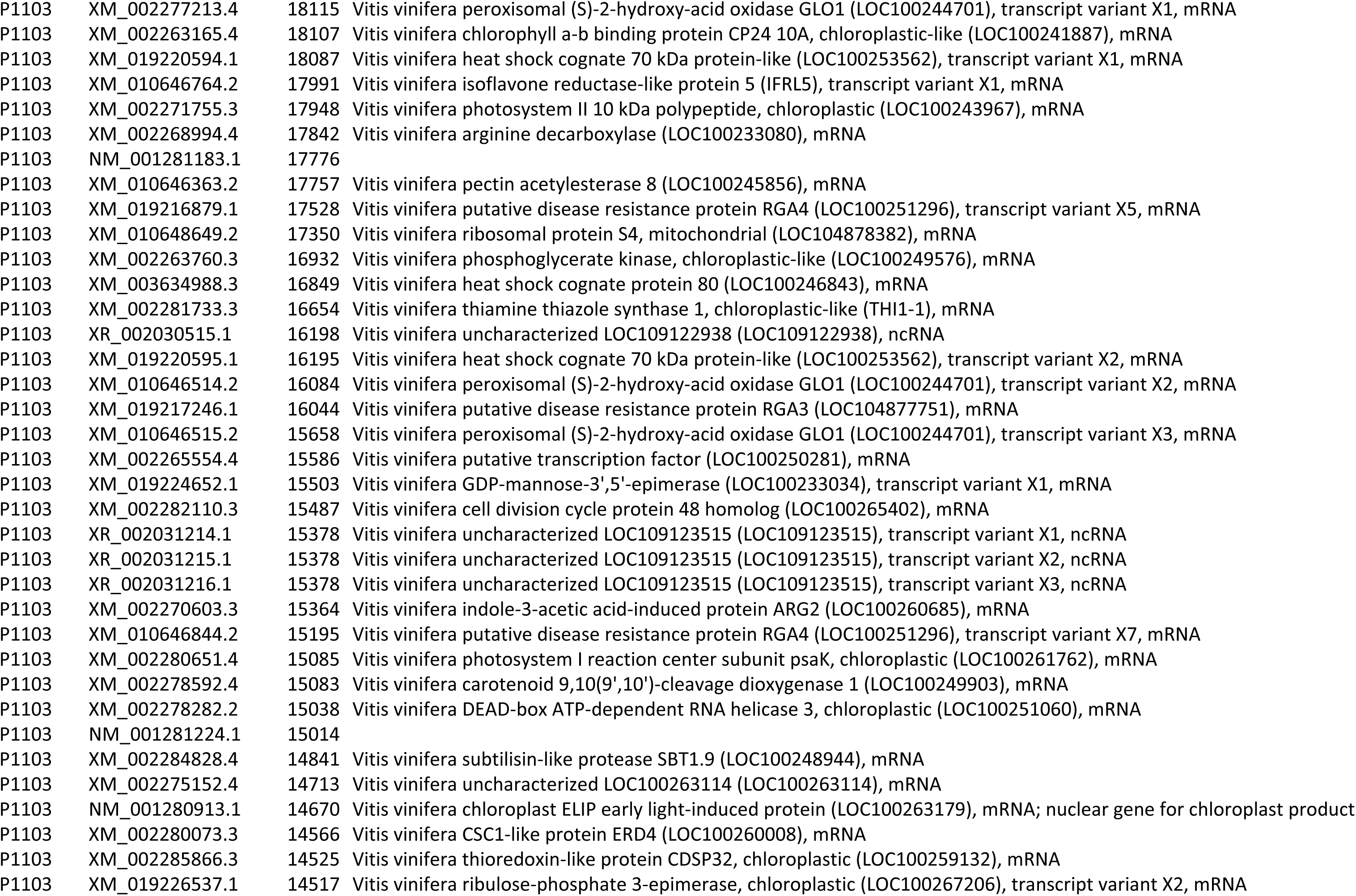
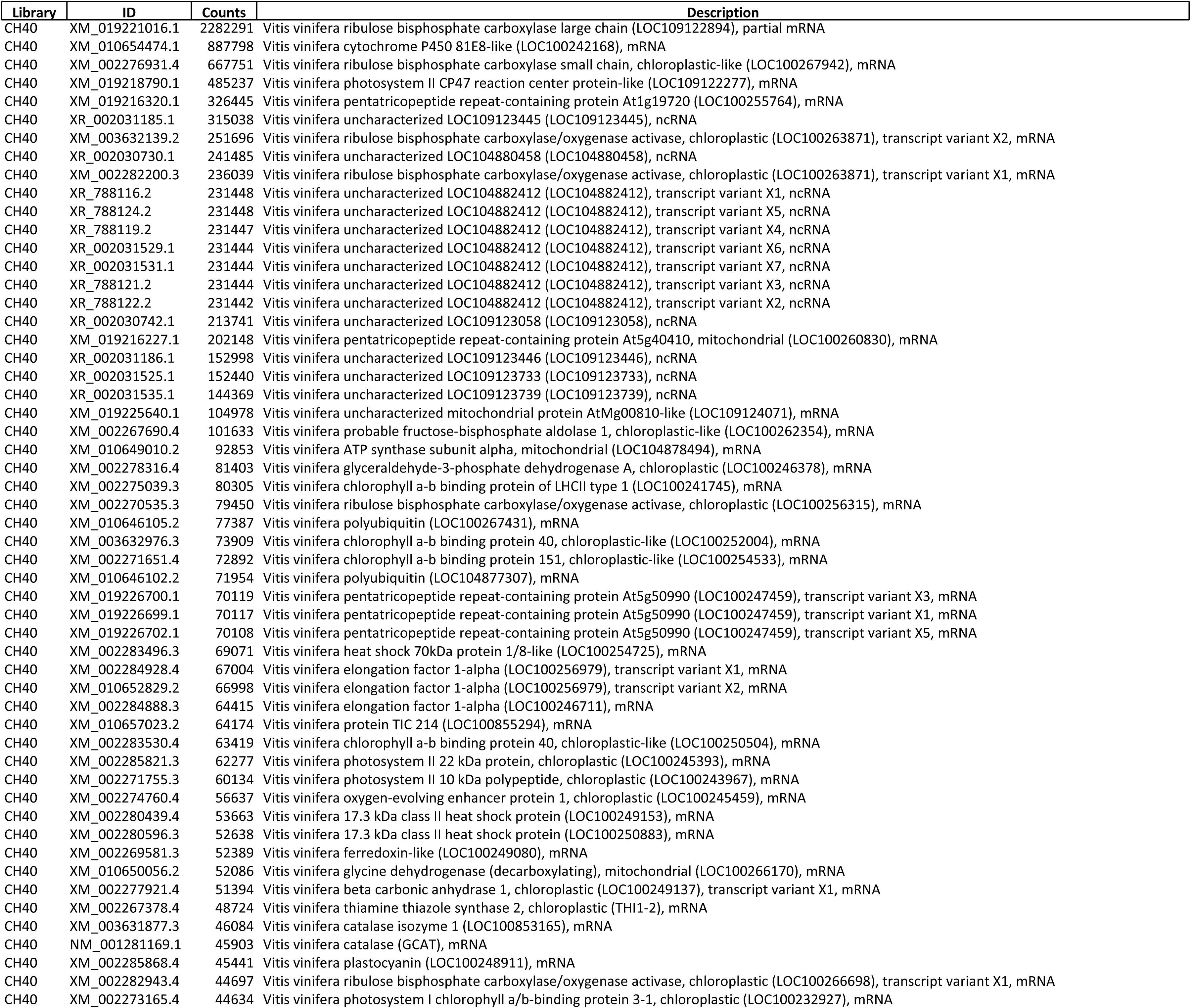
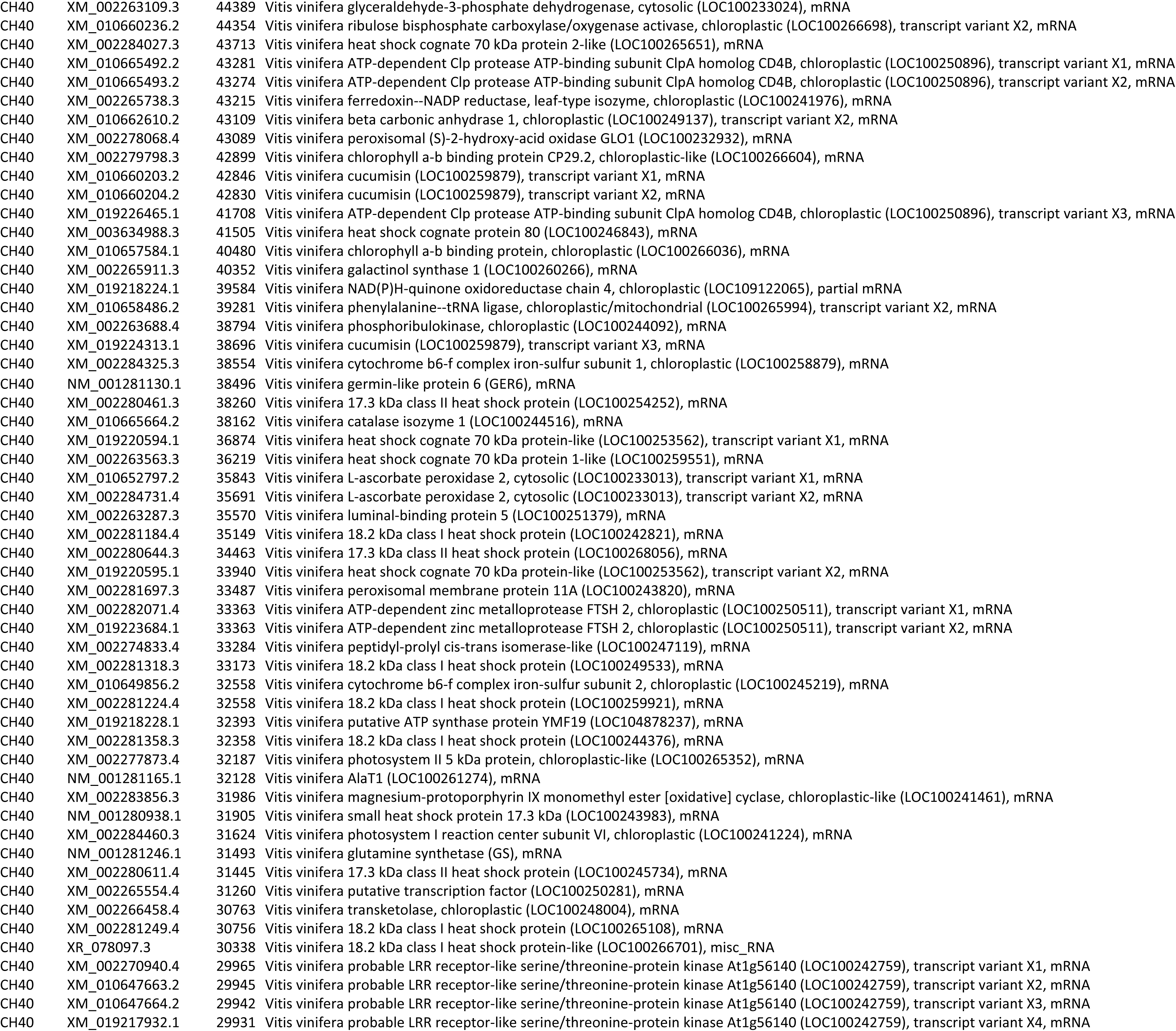
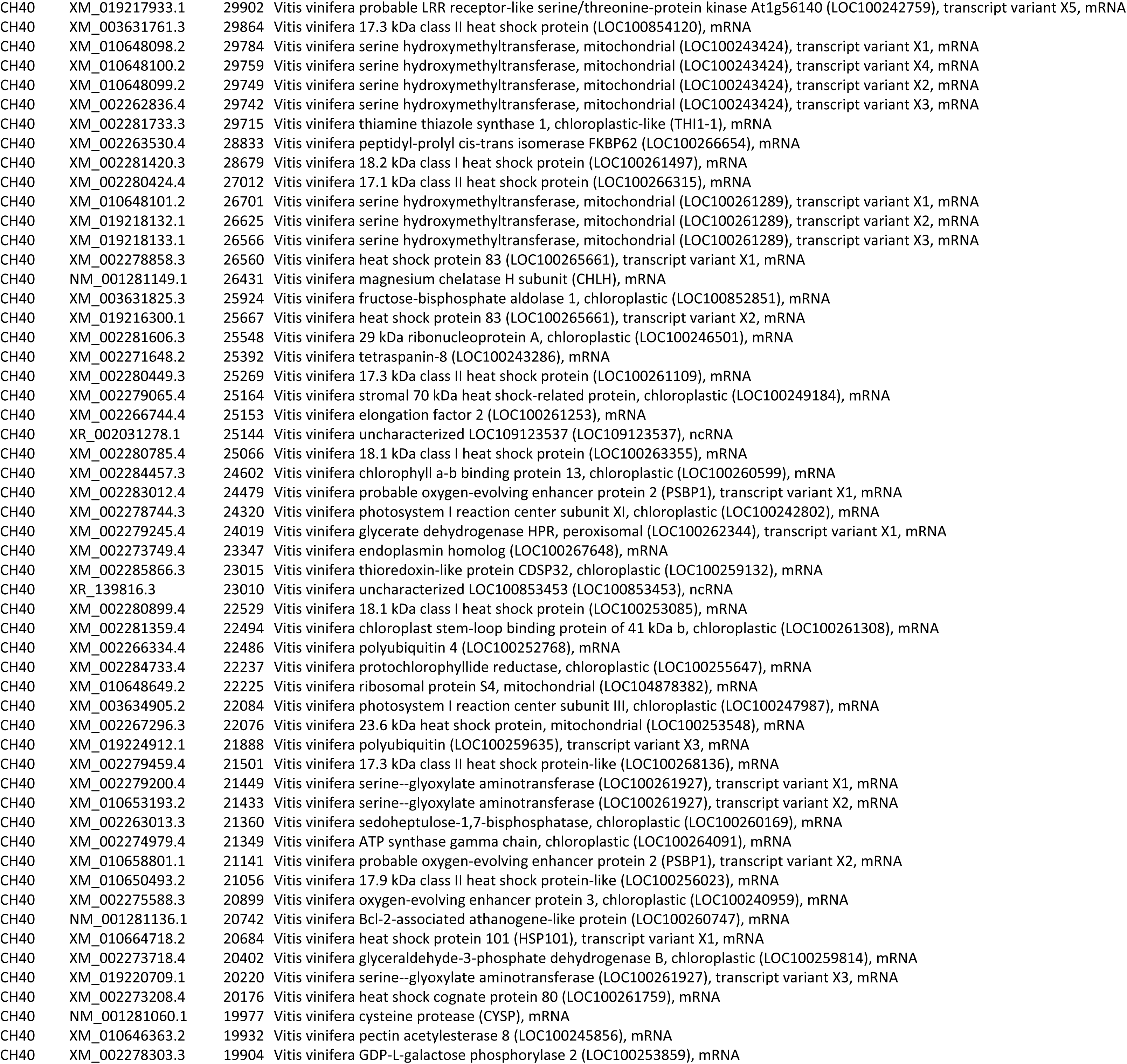
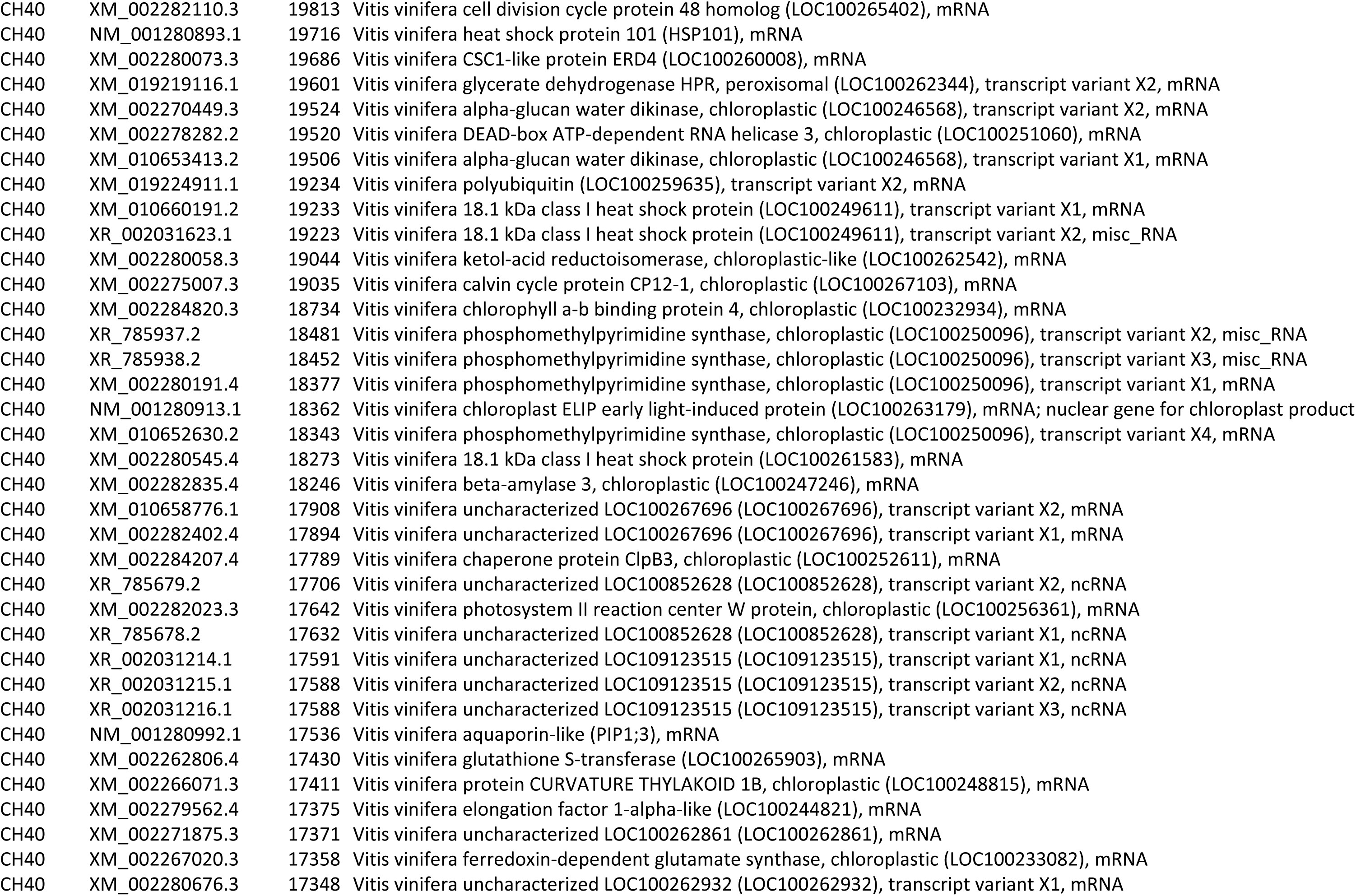
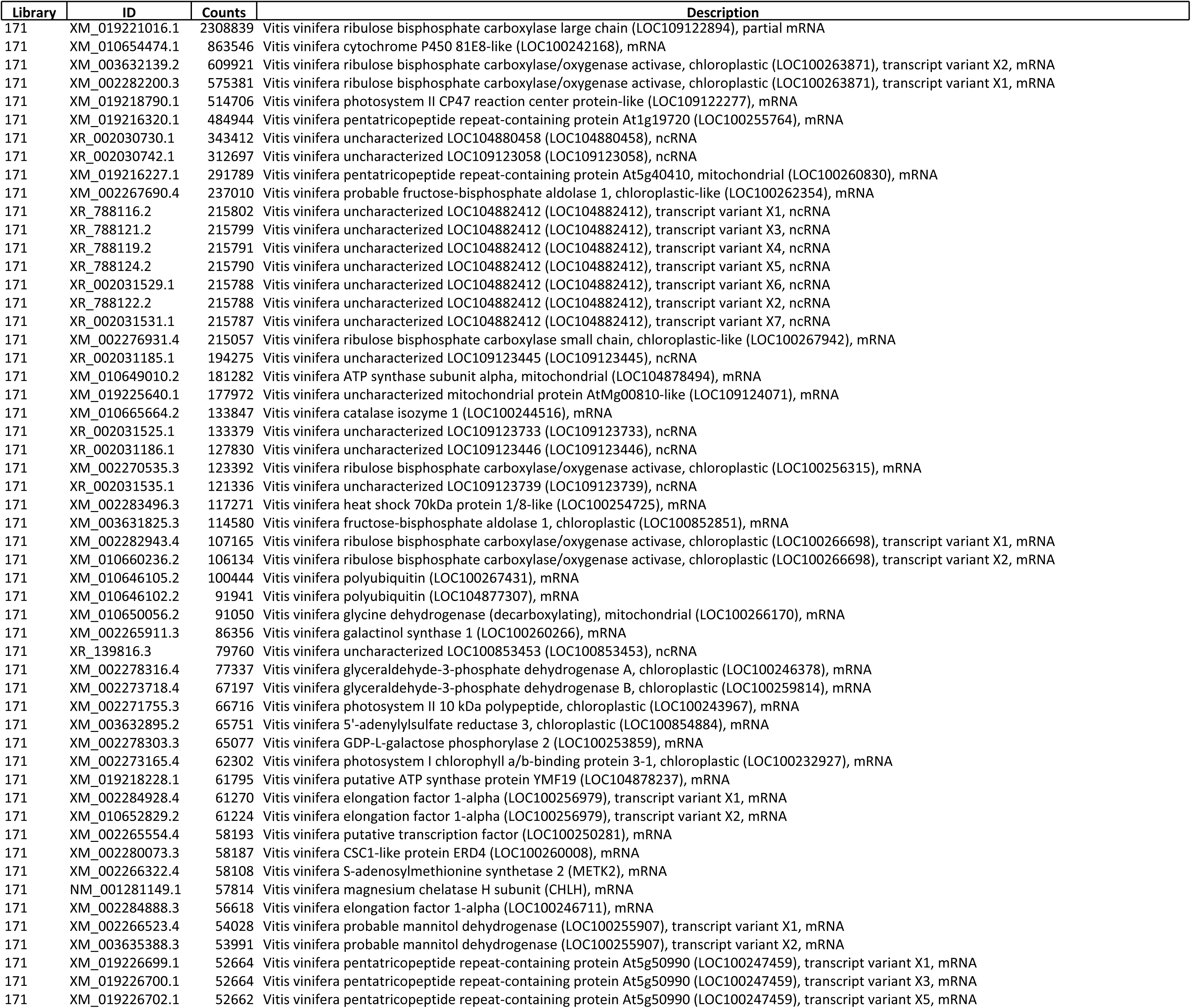
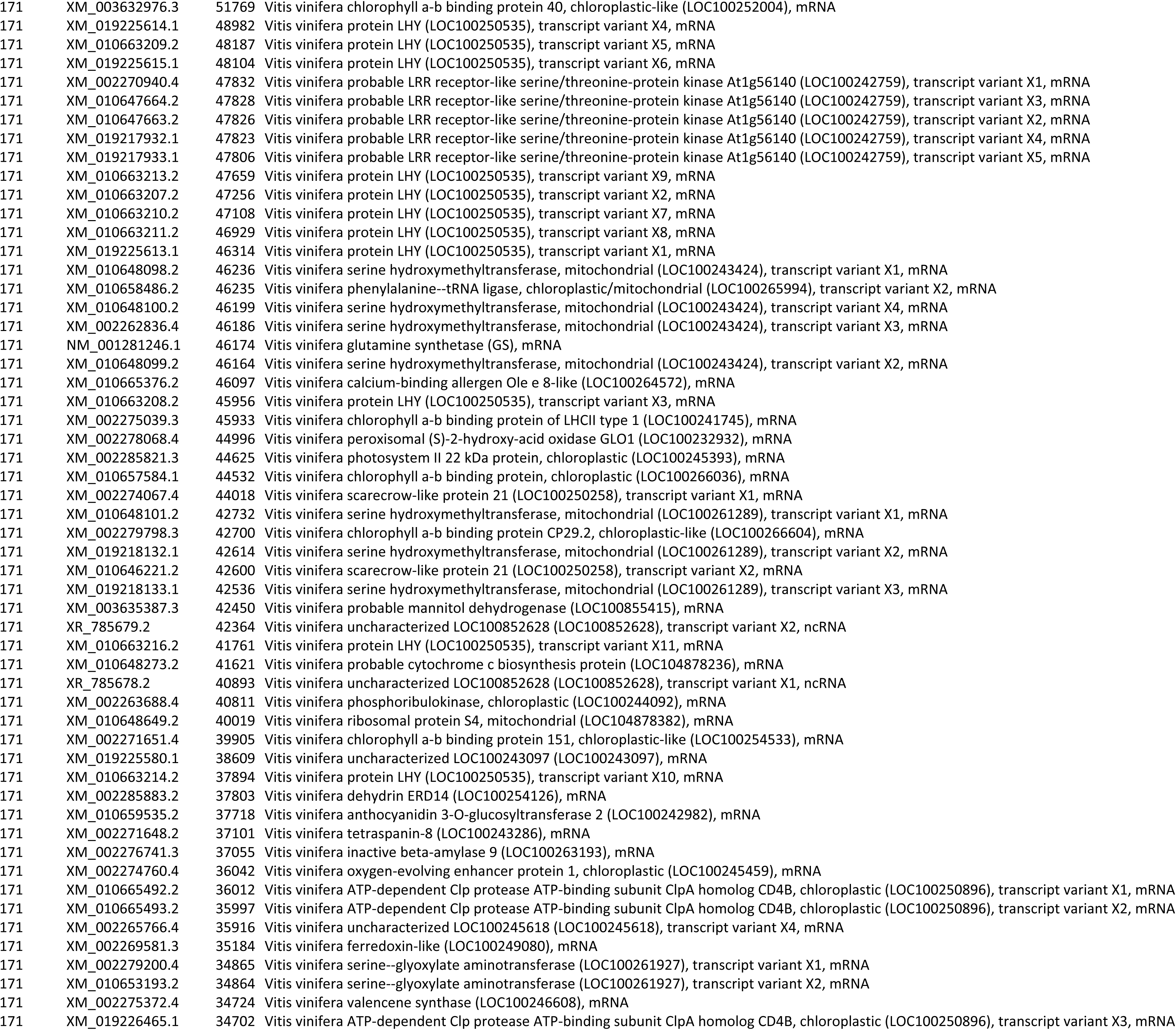
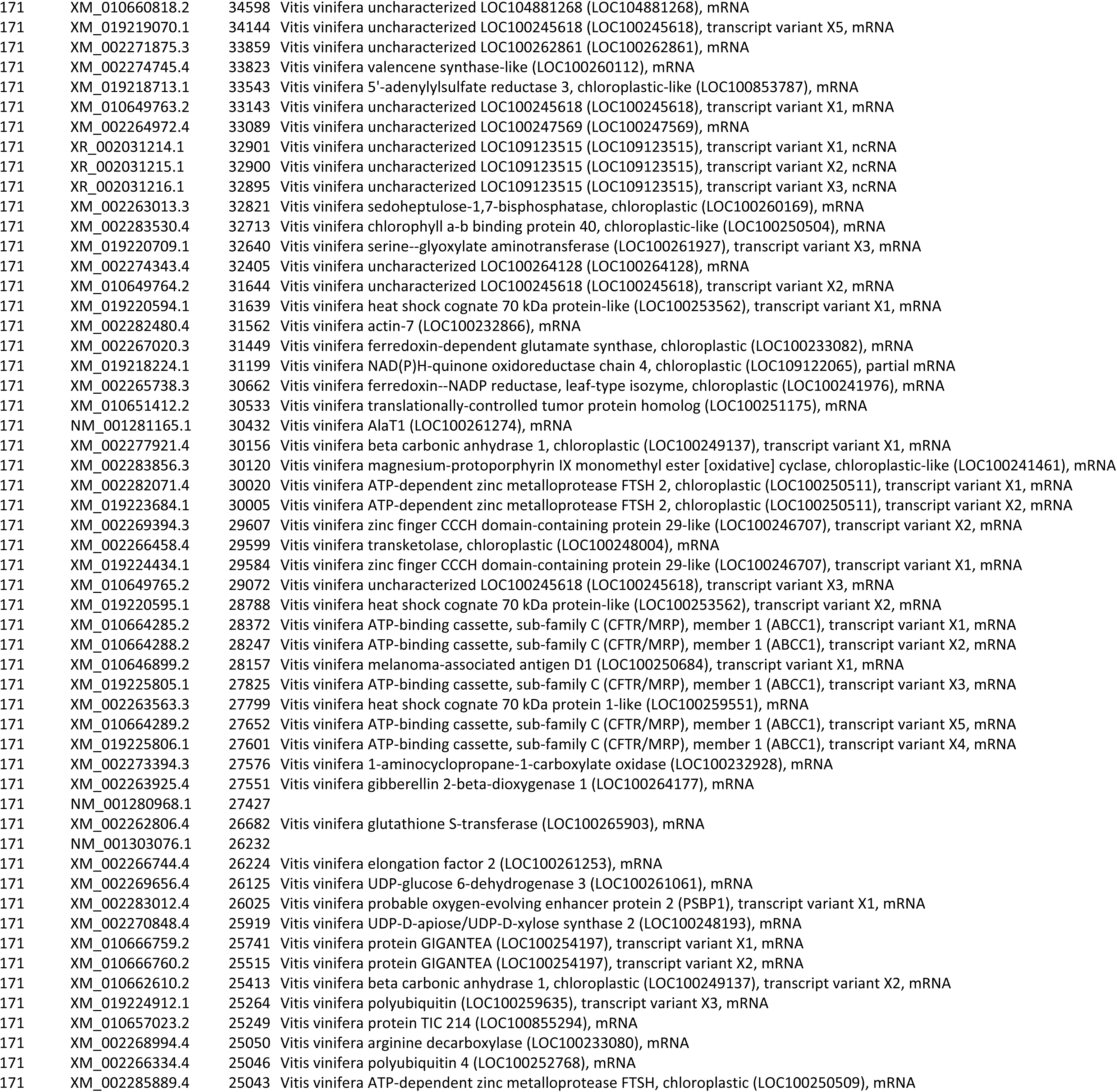
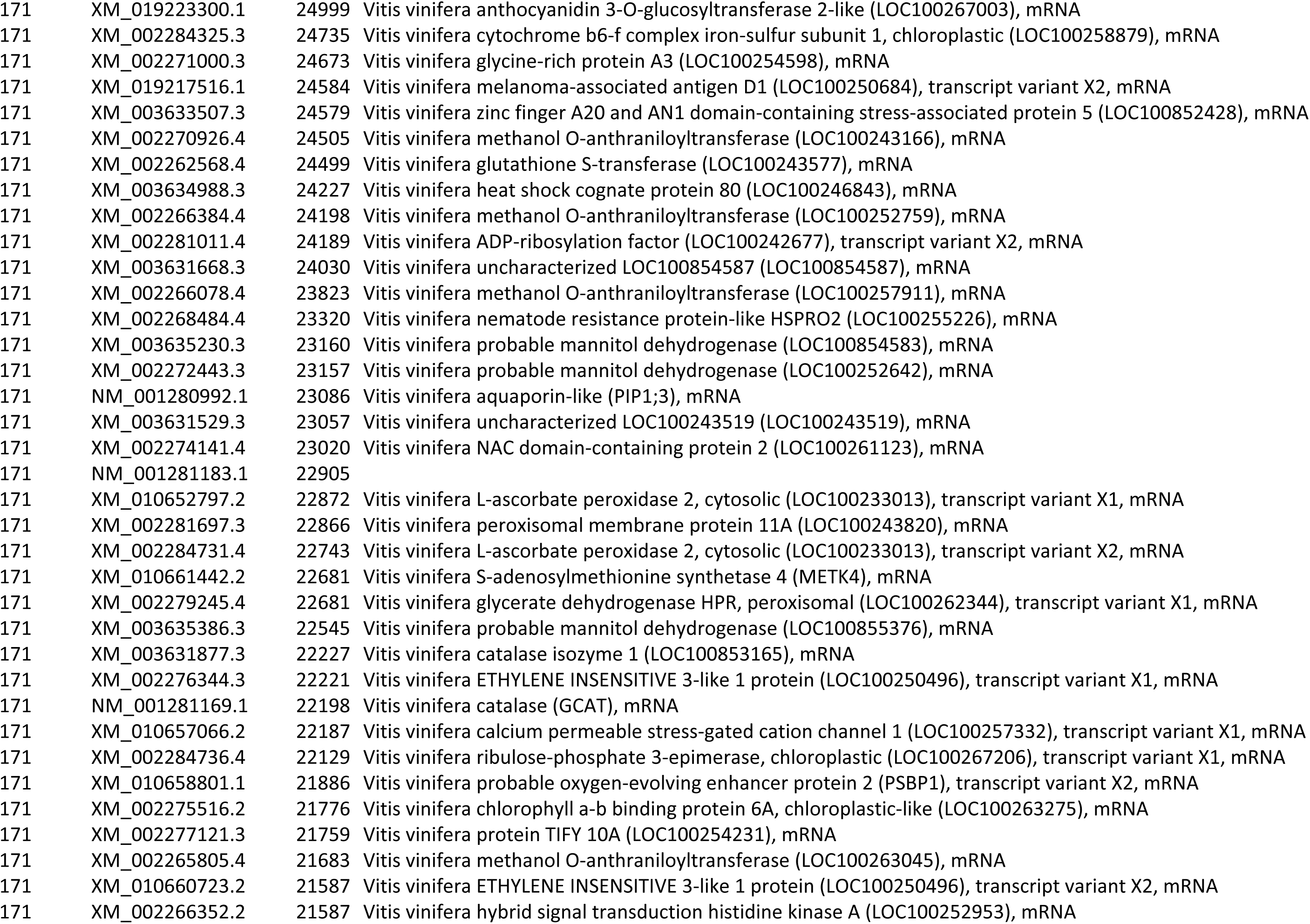
Top 200 most expressed transcripts per library (MB21, P1103, CH40, 171), ranked by counts; IDs and descriptions standardized to RefSeq (XM_/XR_/NM_). Here, NM_ denotes curated RefSeq mRNAs, XM_ predicted model mRNAs, and XR_ predicted model non-coding RNAs. A small subset of NM_ accessions lacks an associated description in this dataset (left blank).

**Supplementary Table 3.** Analysis of the increase in enrichment ratio. Different subtractions comparing an endogenous gene such as actin and different ribosomal genes (18s 1, 2, 3, and 28s listed in Table 3c) against a viral gene such as GLRaV-2 in a 10:1 fold between subtractor and target. Note that although the results for 18s 2 and 28s appear favorable by reducing the expression of ribosomal genes by 3 to 4, even 5 times, they were not reproducible, neither at the same 10:1 ratio nor at a higher 50:1 ratio.

| Genes comparison | Enrichment 10:1 |
| --- | --- |
| ACT vs GLRaV-2 | 0.16706715 |
| 18s 1 vs GLRaV-2 | 0.00280944 |
| 18s 2 vs GLRaV-2 | 4.58683814 |
| 18s 3 vs GLRaV-2 | 0.14890871 |
| 28s vs GLRaV-2 | 3.17663647 |

| Genes comparison | Enrichment 50:1 |
| --- | --- |
| ACT vs GLRaV-2 | 0.8288065 |
| 18s 1 vs GLRaV-2 | 0.01393799 |
| 18s 2 vs GLRaV-2 | 2.75586723 |
| 18s 3 vs GLRaV-2 | 0.738754371 |
| 28s vs GLRaV-2 | 5.759668360 |

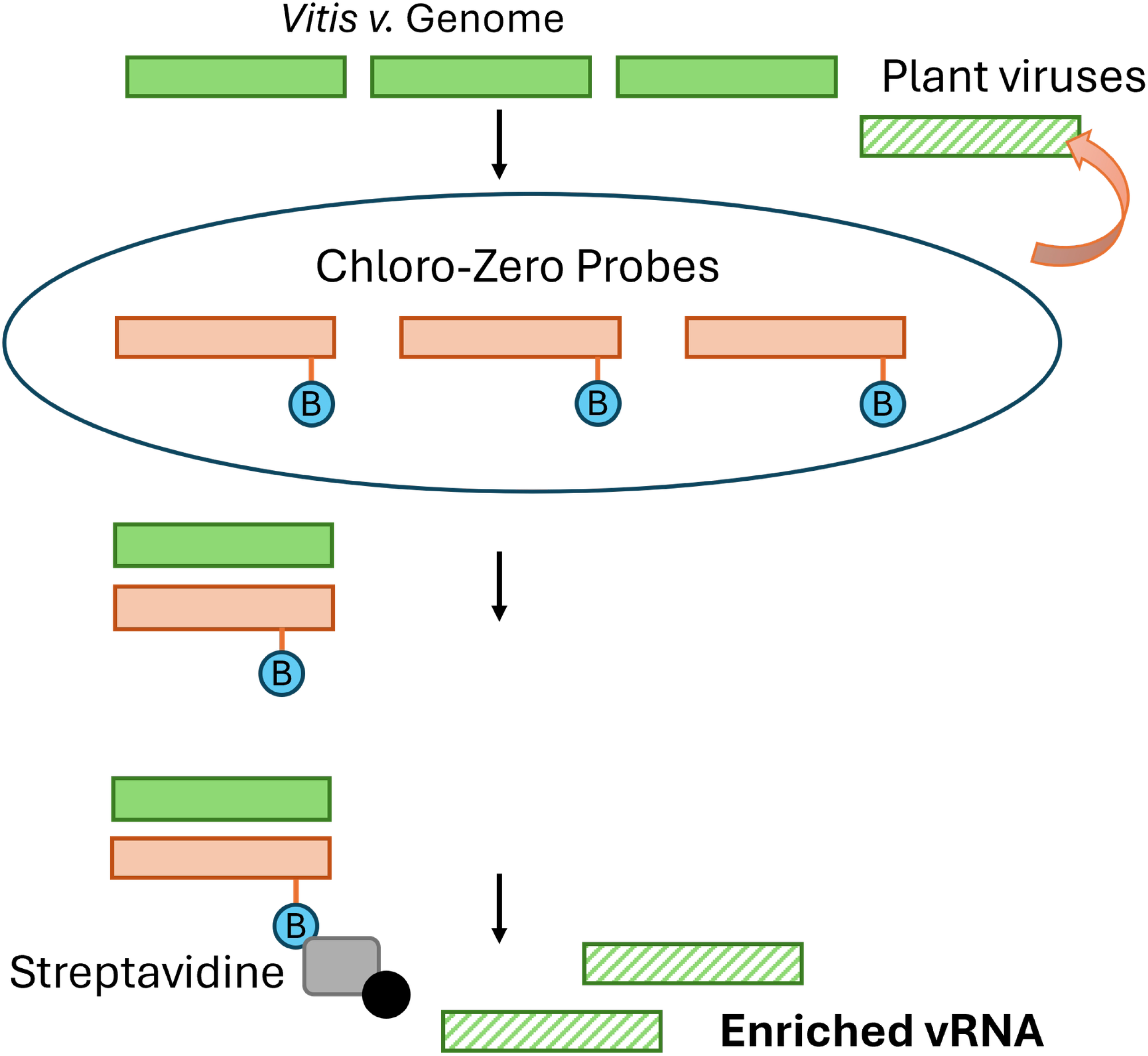

